# Transgenic expression of *Nix* converts genetic females into males and allows automated sex sorting in *Aedes albopictus*

**DOI:** 10.1101/2021.07.28.454191

**Authors:** Célia Lutrat, Roenick P. Olmo, Thierry Baldet, Jérémy Bouyer, Eric Marois

**Author notes:** J.B. and E.M. contributed equally to this work. **Contributions** C.L., T.B., J.B. and E.M. designed research. C.L., R.P.O. and E.M. performed research and analysed data. C.L. wrote the paper with inputs from all authors.

## Abstract

*Aedes albopictus* is a major vector of arboviruses. Better understanding of its sex determination is crucial for developing mosquito control tools, especially genetic sexing strains. In *Aedes aegypti, Nix* is the primary gene responsible for masculinization and *Nix*-expressing genetic females develop into fertile, albeit flightless, males. In *Ae. albopictus, Nix* has also been implicated in masculinization but its role remains to be further characterized.

In this work, we established *Ae. albopictus* transgenic lines ectopically expressing *Nix*. Several were composed exclusively of genetic females, with transgenic individuals being phenotypic and functional males due to the expression of the *Nix* transgene. Their reproductive fitness was marginally impaired, while their flight performance was similar to controls. Overall, our results show that *Nix* is sufficient for full masculinization in *Ae. albopictus*. Moreover, the transgene construct contains a fluorescence marker allowing efficient automated sex sorting. Consequently, such strains constitute valuable sexing strains for genetic control.

---

*Aedes albopictus* can transmit the main human arboviruses including dengue, chikungunya, yellow fever and Zika viruses ^1–6^. Given the global expansion of this mosquito ^7,8^ and its increasing insecticide resistance ^9,10^, new and more sustainable vector control tools are urgently needed ^11^. Genetic control methods to decrease mosquito population, including the Sterile Insect Technique ^12^ and the Incompatible Insect Technique ^13^, have proven effective in several field trials ^14–17^. These methods rely on mass releases of male mosquitoes that compete with their field counterparts for mating. However, in order to upscale the mass production process, an automated and more cost-efficient method for separating males from females is still necessary ^18,19^. Unfortunately, reliable sexing strains are not yet available on an operational scale in *Aedes albopictus*, in part due to the limited knowledge of the sex determination in this species.

Insects exhibit a large diversity of primary signals that activate the sex determination cascade. These signals are poorly conserved between insect genera and few have been discovered so far. In *Drosophila melanogaster*, the chromosome X:autosome ratio is the primary signal for sex determination and dosage compensation. While the Y chromosome is essential for male fertility, it does not determine phenotypic sex ^20^. In some non-Drosophilid Diptera, the primary signal relies on a male factor (M-factor) located either on the Y chromosome or in the M-locus of an autosome. *Aedes* mosquitoes carry an M-factor located within an M-locus on the first pair of autosomes, which is called *Nix* ^21–23^. *Nix* was first identified in *Aedes aegypti*, where it consists of two exons separated by a 99-kb intron, and encodes a 288-amino acid protein ^21^. Stable transgenic expression of *Nix* in *Ae. aegypti* has been shown to be sufficient for converting all genetic females into phenotypic fertile males, confirming its role ^24^. However, these phenotypic males were unable to fly, most likely because they lacked *myo-sex*, another gene closely linked to the M-locus encoding a male-specific flight muscle myosin. In *Ae. albopictus, Nix* is also located on chromosome I and comprises two exons with high homology to *Ae. aegypti Nix* and a much shorter intron ^25^, as well as two additional exons uncovered recently ^22^. *Nix* disruption using CRISPR/Cas9 leads to partial feminization of males, confirming the role of *Nix* as the M-factor in this species ^22^.

It remains to be shown whether *Ae. albopictus Nix* alone is sufficient for masculinization and if genetic females transformed into phenotypic males by transgenic expression of *Nix* would be fertile and able to fly.

In this work, we generated transgenic *Ae. albopictus* mosquitoes expressing the four main *Nix* isoforms in genetic females and showed that all can induce partial to complete masculinization. Some of these lines constitute valuable genetic sexing strains, allowing high-throughput sex sorting at the neonate stage.

## Results

### Obtaining *Nix*-expressing transgenic *Ae. albopictus* lines

Because the other exons were not yet reported when this project began, we first built a transgenesis plasmid comprising *Nix* exons 1 and 2 under the control of the endogenous *Nix* promoter (**Supplementary Data 1**) and an eGFP marker under the control of the OpIE2 promoter. About 300 *Ae. albopictus* embryos were injected, their offspring analysed over three generations and five fluorescent transgenic lines of interest were selected, in which all males were transgenic (for detailed procedures see **Supplementary Note 1**). Lines which yielded non-fluorescent males and non-masculinized eGFP-positive females were discarded. The second injection mix comprised three different transgenesis plasmids designed to express the various *Nix* isoforms ^22^, labelled with different fluorescence markers (eGFP, YFP, and DsRed) under the control of the *Ae. aegypti* poly-ubiquitin promoter (PUb). We injected about 700 embryos with this mix and selected seven transgenic lines of interest (see **Supplementary Note 1**).

### Identification of transgenic lines devoid of an M-locus

During characterization of *Nix*-expressing transgenic lines, we observed strong genetic linkage between fluorescence and the male phenotype. This could be due either to transgenic fluorescent males actually being masculinised genetic females, resulting in mosquito lines lacking natural males; or to the insertion of the piggyBac plasmid cassette within or near the endogenous M-locus, resulting in genetic linkage between the fluorescent transgene and male sex. To distinguish between these two possibilities, we tested for the presence of endogenous and exogenous *Nix* by PCR (e.g. see **Supplementary Figure 1**). In eight out of twelve lines, endogenous *Nix* was detected in the GFP positive transgenic males. Therefore, in these lines, maleness was natural and the transgene was M-linked. In contrast, four of our mosquito lines were devoid of endogenous *Nix*, namely SM9, 1.2G, 2.2G and 3.1G. Interestingly, these four lines possessed the eGFP fluorescent marker, thus, they expressed the shortest *Nix* isoforms encoded by exons 1 and 2 only (**Supplementary Figure 2**). In some lines marked by YFP and/or DsRed fluorescence, a fraction of the males lacked the M-locus, while others possessed it. However, the M-deprived males did not sire any progeny, thus they were possibly sterile.

Single males from the four M-free GFP lines were backcrossed to negative females for several generations to eliminate additional non-fully masculinizing transgene insertions (e.g. see **Supplementary Figure 3**). Following this step, we obtained three lines (SM9, 1.2G and 3.1G) where only males showed eGFP fluorescence, without residual fluorescent females or intersex individuals. Further tests and analyses were performed on the SM9 line, and some experiments were replicated on the other two lines.

### Confirmation of the role of *Nix* transgenes in masculinization

To further confirm the role of *Nix* transgenes in masculinization, we injected SM9 embryos with a plasmid expressing CRE recombinase in order to excise the *Nix*-eGFP transgenic cassette, which is flanked by lox recombination sites (**Supplementary Figure 2**). At the pupal stage, efficiently injected individuals showed a striking loss of eGFP expression at the injection site (posterior pole), indicative of lox cassette excision. While these pupae should have developed into phenotypic males (due to the eGFP-marked *Nix* transgene cassette), they showed female genitalia, hence demasculinization of their posterior pole (**Figure 1A-C, Supplementary Figure 4**). Adult mosquitoes hatching from these pupae showed antero-posterior gynandromorphism, having male heads and female genitalia. Strikingly, a single individual, which arose from an embryo accidentally injected in the anterior pole rather than in the posterior, showed the opposite gyndandromorphic phenotype, with a female head and male genitalia (**Figure 1D**). These results confirmed that maleness in the SM9 line results from the transgene’s activity, which can be abolished by CRE/lox excision. The co-existence of male and female tissues in the same individual also illustrates that sex determination is tissue-autonomous in *Ae. albopictus*.

**Figure 1:**
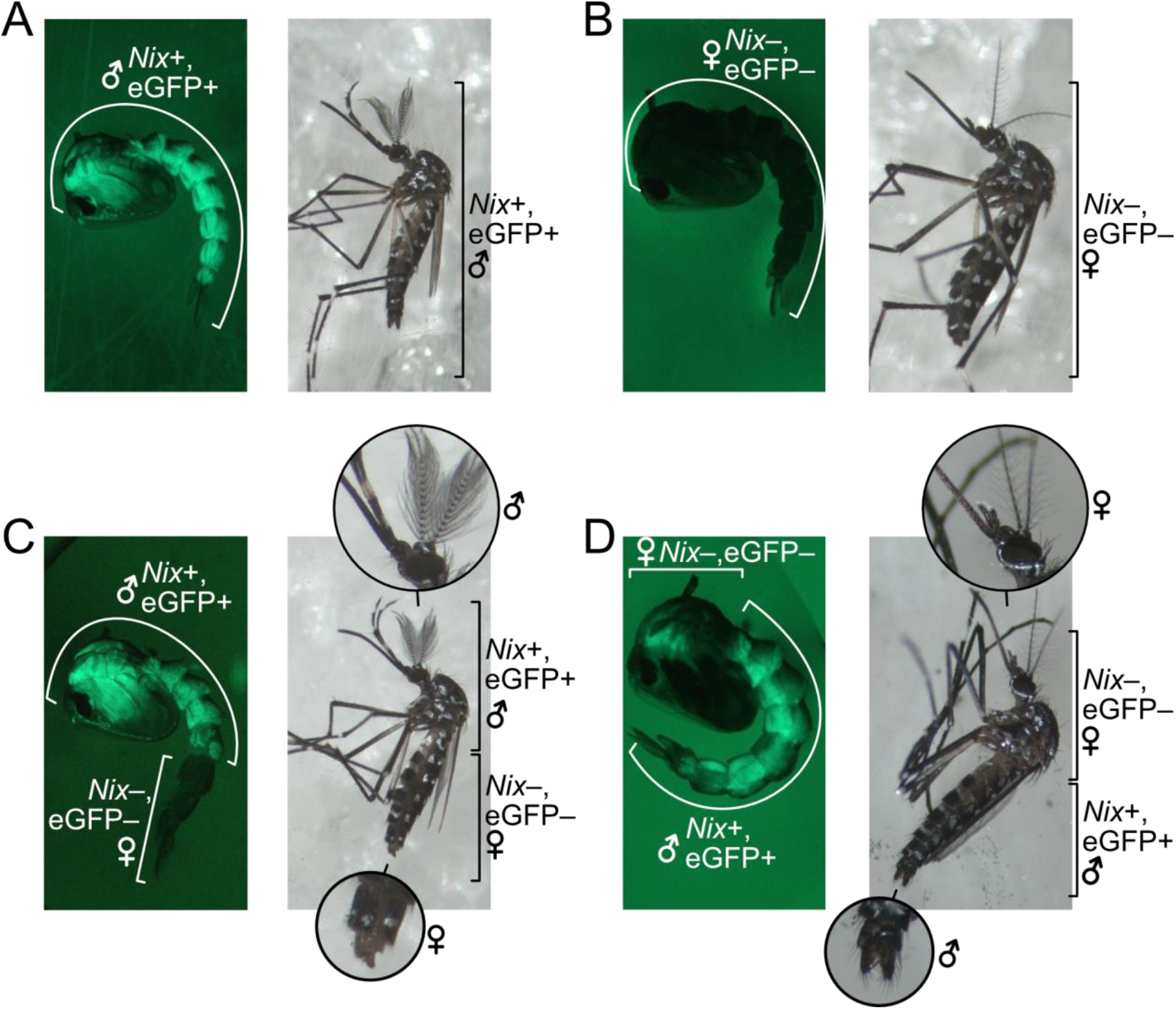
Tissue demasculinization upon CRE/lox excision of the transgenic *Nix* cassette. **A)** Representative transgenic male pupa and male adult from the SM9 line. **B)** Representative non-transgenic female pupa and female adult from the SM9 line. **C)** Transgenic SM9 male pupa and adult injected as embryos in the posterior pole with CRE-recombinase that excised the Nix-eGFP cassette in the injected region. These individuals show a male anterior body with female genitalia. **D)** Transgenic SM9 male pupa and adult injected with CRE-recombinase in the anterior pole of the embryo. Note the female anterior body and male genitalia.

### Characterization of *Nix*-expressing m/m pseudo-males

To evaluate whether the *Nix-eGFP* cassette was stable and adult pseudo-males fully viable, we first determined the sex ratio in comparison to that of the parental wild strain (WT). Male ratios from SM9 (sex-ratio estimate ± SE = 0.56 ± 0.05), 1.2G (0.49 ± 0.05) and 3.1G (0.54 ± 0.05) transgenic strains were not significantly different from that of the WT line (0.52 ± 0.10, SM9 *vs*. WT *p*-value = 0.297, 1.2G *vs*. WT *p*-value = 0.558, 3.1G *vs*. WT *p*-value = 0.869).

In *Aedes* mosquitoes, males and females display a significant size dimorphism, with females having a larger body size ^26^. We determined the body size of pseudo-males using wing length as a proxy ^27^. Our results showed no significant difference between the size of the wild-type and transgenic males (WT male *vs*. SM9 pseudo-male p = 0.998), while both were significantly different from females (WT male *vs*. WT female *p*-value < 0.001, SM9 pseudo-male *vs*. WT female *p*-value < 0.001, **Figure 2**).

**Figure 2:**
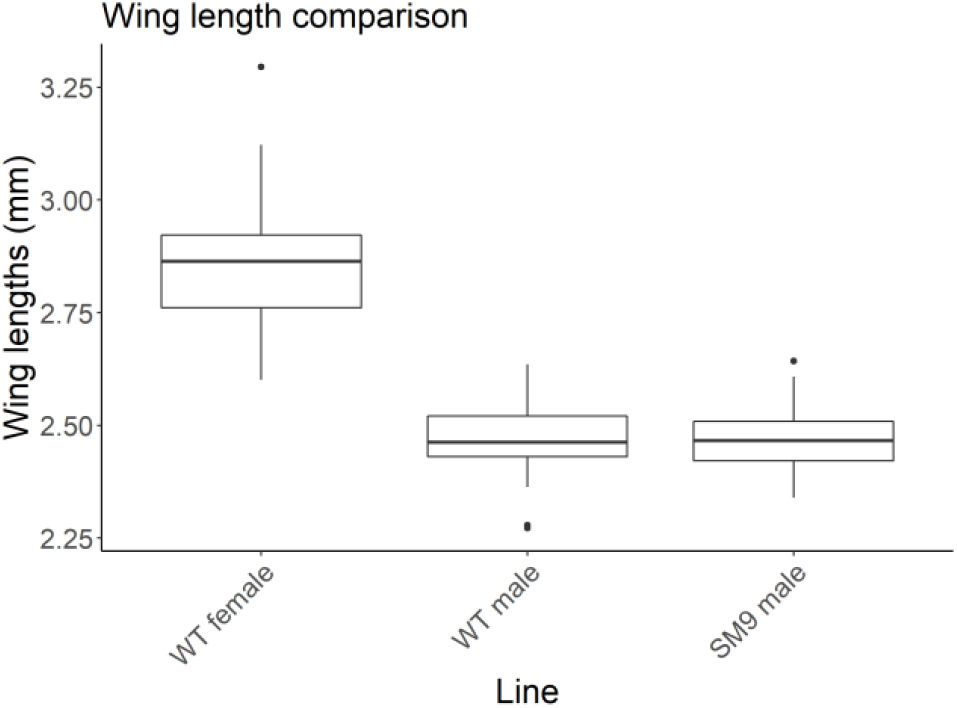
Wing length comparison between wild-type *Ae. albopictus* males and females and *Nix*-expressing SM9 pseudo-males. Wing length was measured on ImageJ software from pictures of dissected right wings taken under a binocular microscope. WT male *vs*. WT female *p*-value < 0.001, SM9 male *vs*. WT female *p*-value < 0.001, WT male *vs*. SM9 male *p*-value = 0.998.

*Nix* expression levels in pseudo-males *vs*. wild-type males were compared by RT-qPCR at the pupal stage (**Figure 3A**). Interestingly, pseudo-males from three different lines expressed *Nix* at a similar level to wild-type males (SM9 males *vs*. WT males *p*-value = 0.501, 1.2G males *vs*. WT males *p*-value = 0.391, 3.1G males *vs*. WT males *p*-value = 0.626). Thus, we inquired if the downstream double switch genes, *doublesex* (*dsx*) and *fruitless* (*fru*), showed male-specific splicing products in pseudo-males. For this, we performed RT-PCR on pseudo-males of four transgenic lines. Results revealed that pseudo-males displayed the same splicing pattern as WT males for both genes (**Supplementary Figure 5**).

**Figure 3:**
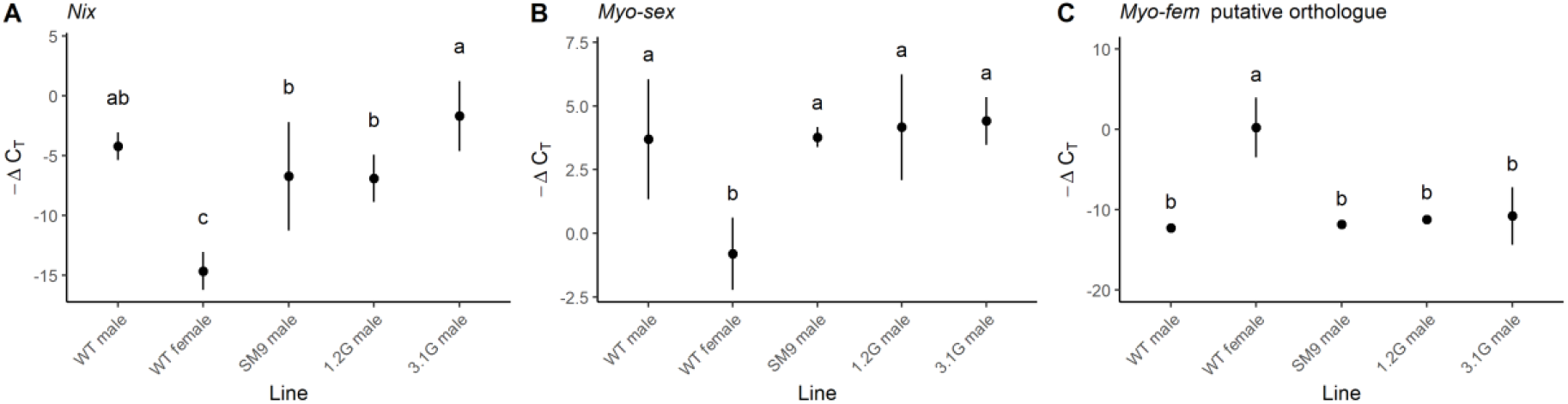
Relative expression of *Nix, myo-sex* and *myo-fem* orthologue in *Ae. albopictus* transgenic males. RT**-**qPCR results are represented by -ΔC_T_, which reflects the relative expression level of each gene in a given treatment, C_T_ values being inversely proportional to the expression levels. *AalRps7* was used as endogenous reference gene. On each panel, distinct letters represent significant difference in pairwise Tukey test (*p* < 0.001). Dots represent the mean value of the 3 biological replicates, vertical lines represent 95% confidence interval (CI). **A)** *Nix* relative expression. **B)** *myo-sex* relative expression. **C)** Relative expression of the candidate orthologue of the *Ae. aegypti myo-fem* gene, LOC109402113.

### Flight ability of Nix-expressing pseudo-males

In contrast to what has been reported in *Ae. aegypti* where the lack of the M-linked gene *myo-sex* resulted in flightless pseudo-males ^24^, our *Ae. albopictus* pseudo-males were readily able to fly. Consequently, we compared the expression levels of the *Ae. albopictus* orthologues of the sex-specific flight muscle myosins identified in *Ae. aegypti, myo-sex* and *myo-fem*, in *Nix*-expressing pseudo-males *vs*. WT males by RT-qPCR. We discovered an M-linked copy of the *myo-sex* gene (**Supplementary Data 2**), additional to those already known ^28^, as well as an orthologue of *myo-fem* (**Supplementary Note 2**). Our results showed that the collective level of *myo-sex* expression (all copies have 100% cDNA identity) in WT females was approximately 20 fold lower compared to males (*p*-values < 0.001), and that pseudo-males expressed *myo-sex* at a level similar to wild type males (SM9 males *vs*. WT males *p*-value = 1.000, 1.2G males *vs*. WT males *p*-value = 0.960, 3.1G males *vs*. WT males *p*-value = 0.843, **Figure 3B**). These results suggest that one or several non M-linked, endogenous *myo-sex*-like copies are efficiently upregulated in pseudo-males. Additionally, pseudo-males express *myo-fem* at similarly low levels as wild-type males (SM9 males *vs*. WT males *p*-value = 0.995, 1.2G males *vs*. WT males *p*-value = 0.841, 3.1G males *vs*. WT males *p*-value = 0.614, **Figure 3C**). Gene expression in all males tested was approximately 10,000 fold lower than in wild-type females (*p*-values < 0.001 for all combinations, **Figure 3C**).

Since we observed that genes potentially involved in flight are regulated similarly in *Nix*-expressing pseudo-males comparing to wild-type counterparts, we tested the SM9 pseudo-males’ flight ability by performing a flight test as described in ^29^. We observed that SM9 males had a higher escape probability than WT males (p-value < 0.001, **Figure 4A**), suggesting that the flight capacity of SM9 pseudo-males was at least as high as that of WT male mosquitoes.

**Figure 4:**
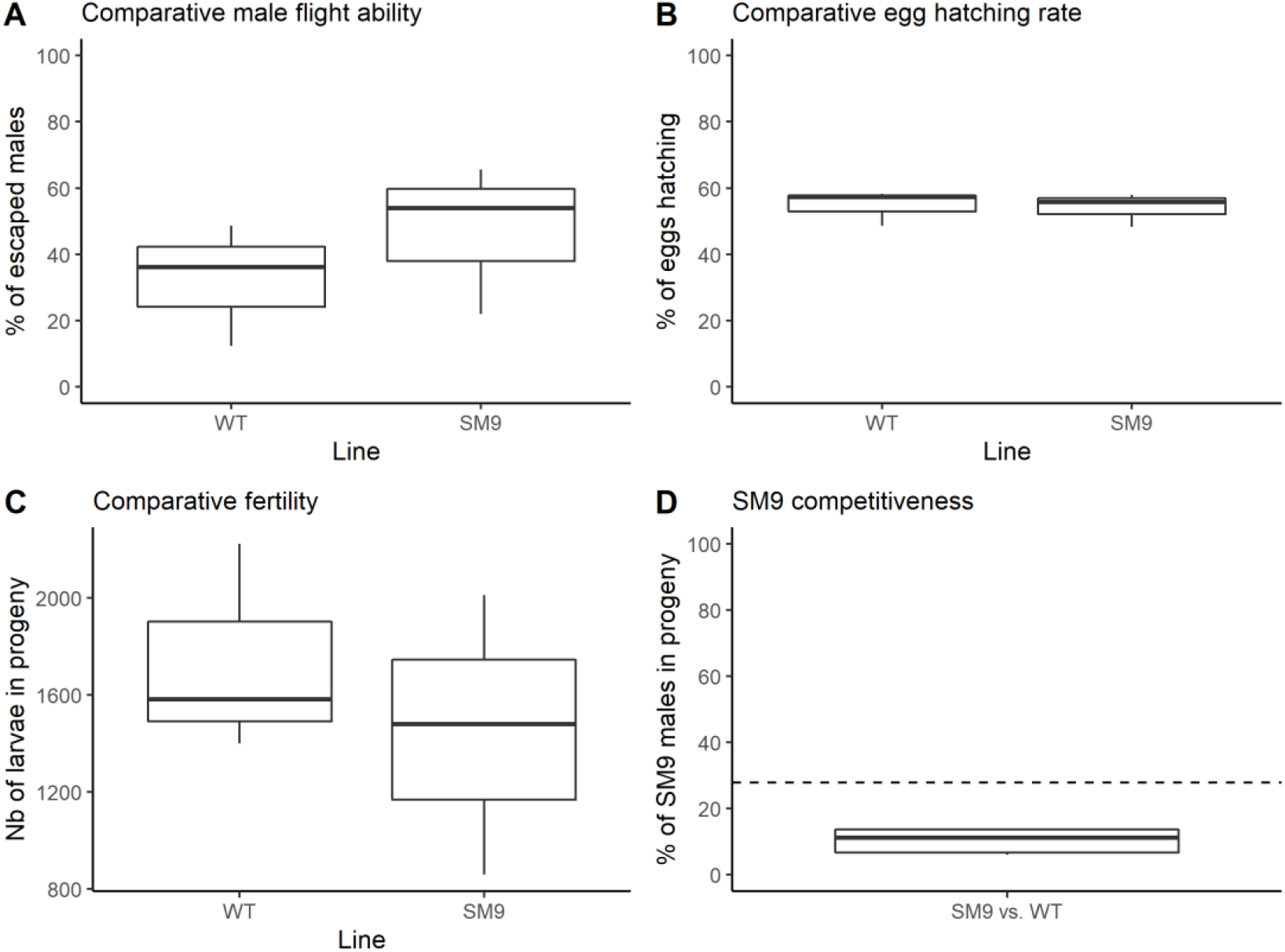
Fitness comparison of *Ae. albopictus* SM9 versus WT males.was measured on ImageJ. **A)** Percentage of males that successfully escaped the flight test device. 3 replicates with an average of 82 ± 13 males were performed (N = 489). To test the effect of the lines on the flight test success, we used linear generalised mixed-effect model and Bernoulli distribution assumptions with “replicate” as random effect. *p*-value < 0.001. **B)** Fertility measured by the number of progeny sired by 30 males crossed with 60 females (replicated 3 times per line). *p*-value = 0.532. **C)** Hatching rate measured by dividing the number of progeny by the number of eggs on sub-samples of the egg batches. Dried eggs were counted, submerged, placed in a vacuum chamber for 30mn and allowed to hatch for 24h before counting larvae (replicated 3 times per line). *p*-value = 0.423. This data allows the estimation of fecundity, which is the total number of eggs laid. **D)** To estimate SM9 male competitiveness, 5 competition assays were performed, crossing 30 WT males and 30 SM9 males to 30 females. In their progeny, the percentage of NOE-SM9 sons was measured by COPAS and compared to the expected percentage (dashed line). 10.2% (± 0.4 %) of transgenic progeny *vs*. 27.9% of expected value (*p*-value < 0.001).

### Reproductive fitness of SM9 pseudo-males

We compared SM9 to WT relative fertility (total number of live larvae in a given progeny) and fecundity (total number of eggs laid, estimated by dividing the total number of larvae by the hatching rate). We found no significant difference in SM9 hatching rate (*p-*value = 0.423, **Figure 4B**) or fertility (*p*-value = 0.532, **Figure 4C**) comparing to wild-type control, thus no difference in fecundity either. Then, we measured relative competitiveness between SM9 pseudo-males and wild-type males by mixing equal numbers of transgenic and wild-type males with wild-type females. Fecundity and fertility being similar, if both lines were equally competitive, the expected percentage of SM9 males (fluorescent larvae) would be half the percentage of males in the SM9 strain, i.e. 27.9%. In this experiment, we observed an estimated mean of 10.2 ± 0.4 % SM9 male progeny indicating a reduced competitiveness (*p*-value < 0.001, **Figure 4D**). The same competitiveness assay was performed between 1.2G pseudo-males and WT males and gave a similar result (12.5 ± 0.3 % of transgenic progeny, *p*-value < 0.001).

### Automated sex sorting of transgenic pseudo-males

The transgenesis plasmids carrying a fluorescent marker under strong promoters, neonate larvae can be sorted according to their fluorescence, which, in this case, is sex specific. Similarly to what has been developed in *Anopheles* mosquitoes ^30,31^, sex sorting can be automated using a COPAS device (Union Biometrica) which allows separation of particles based on fluorescence levels. Using COPAS on the SM9 line, we were able to separate green fluorescent males from non-fluorescent females (**Figure 5A**). However, due to the weak activity of the OpIE2 promoter driving GFP at the neonate stage, the fluorescence was not always strong enough to get well separated positive *vs*. negative clouds by COPAS. Lines 3.1G and 1.2G express higher levels of GFP due to the high activity of the PUb promoter at the neonate stage and provide clearer sex separation (**Figure 5B-C**). In all cases, batches of several thousands of neonate larvae could be repeatedly sex separated using COPAS at a speed of approximately 2,400 larvae per minute. Visual screening at the pupal stage confirmed perfect sex separation.

**Figure 5:**
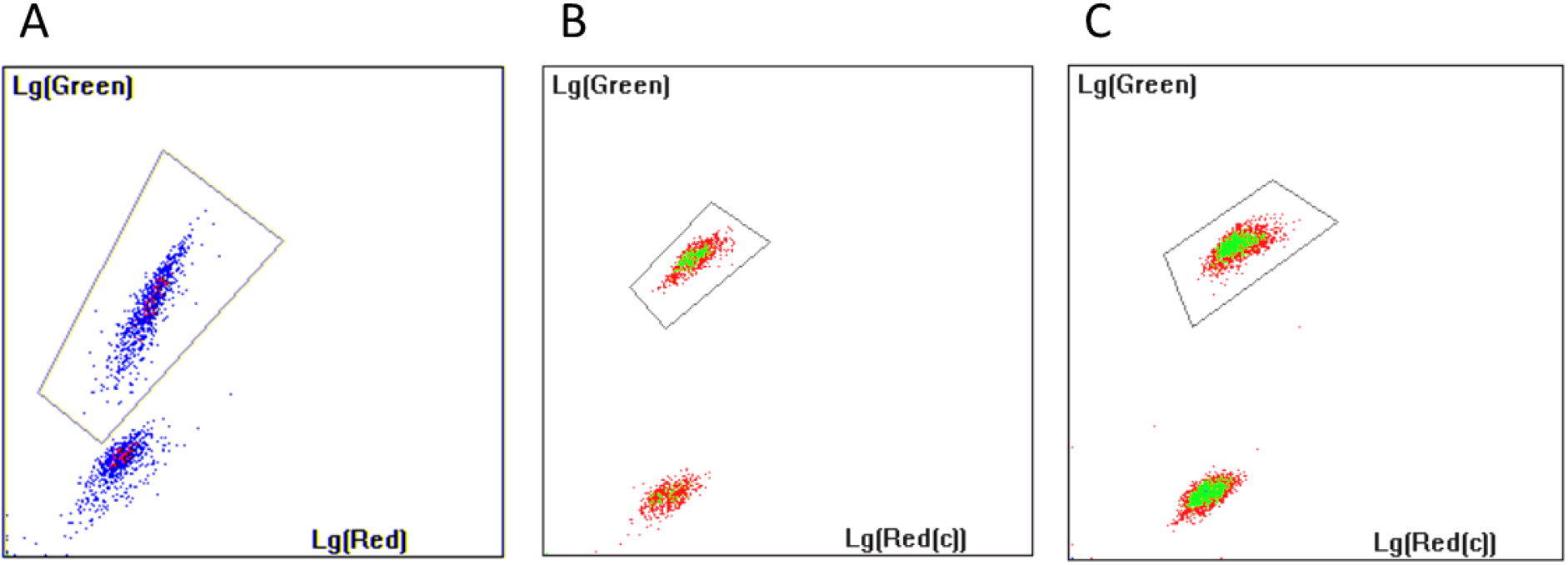
Automated sex sorting of *Ae. albopictus Nix*-expressing transgenic lines. Sorting is performed at the neonate stage using a COPAS device. Circled regions are the ones to be sorted by the machine. Presented graphs allow sorting of larvae based on their red and green fluorescence. *Nix*-expressing pseudo-males being tagged with an eGFP marker gene, the circled region sorts males. **A)** Sorting of 2,246 larvae from the SM9 line. **B)** Sorting of 1,624 larvae from the 1.2G line. **C)** Sorting of 3,876 larvae from the 3.1G line.

## Discussion

The *Nix*-expressing transgenic *Ae. albopictus* lines presented in this work confirm that *Nix* is necessary and sufficient to initiate the male sex determination cascade in *Ae. albopictus*. They also demonstrate that a short version of *Nix* comprising only the first two exons and the first intron is sufficient for complete masculinization. Pseudo-males resulting from the expression of a *Nix* transgene in a female genomic background seem as fecund and fertile as wild-type males.

Similarly to *Ae. aegypti* ^24^, we observed that pseudo-males harbour molecular characteristics of wild-type males. Of note, we were able to detect an undescribed M-linked homologue of *myo-sex* in the *Ae. albopictus* genome, besides additional autosomal copies, characterized by a 600 nucleotide deletion within the gene’s non-coding region. *myo-sex* has been described as responsible for male flight ability in *Ae. aegypti* and is specifically present in wild-type males, hence linked to the M-locus in this species ^24^. However, in *Ae. albopictus*, the M-linked copy does not appear to be essential for male flight, as synthetic males expressing a *Nix* transgene but devoid of the M-linked *myo-sex* copy showed similar total *myo-sex* expression level and performed better in flight tests. Moreover, we also identified a strong candidate for a *myo-fem* orthologue, a gene described in *Ae. aegypti* as essential for female flight ^32^. In *Ae. albopictus*, we measured that this gene is highly expressed in wild-type females and strongly repressed in wild-type males as well as transgenic pseudo-males.

Interestingly, despite good flight ability, SM9 pseudo-males suffered from reduced mating competitiveness. This may result from a negative effect of the ubiquitous expression of the GFP marker gene, from disruption of a gene at the transgene insertion site, or from the absence of unknown factors encoded by the endogenous M-locus involved in male mating ability. Notably, it was observed in *Ae. aegypti* that several long non-coding RNAs of the M-locus were upregulated in the testes and in the male accessory glands ^24,33^. We cannot exclude that similar features are present in the *Ae. albopictus* M-locus and that their absence might influence some aspects of the mating process that we were unable to detect in our fertility and fecundity assays.

Besides fully masculinizing *Nix* transgene insertions, we also obtained a number of insertions triggering no or only partial masculinization, whereas the associated fluorescence reporter genes were expressed under control of the polyubiquitin promoter at comparable levels for all obtained lines. This suggests that the genomic context where the *Nix* transgene is inserted affects its expression, and thus, masculinization. There might be a dose-dependency of *Nix* expression during development with a threshold ensuring full masculinization. However, while this might be true for GFP-bearing cassettes, we failed to isolate a masculinizing line carrying a YFP or a dsRed reporter, (i.e., expressing the long *Nix* isoforms 1 or 2). With these isoforms, we achieved partial to full apparent masculinization, but these individuals were apparently infertile. Therefore, it seems that longer isoforms might not be sufficient for masculinization. Alternatively, the short intron retained in our isoform 3-4 construct (marked by eGFP), which we did not include in the longer isoform constructs, could carry a regulatory sequence that is essential for complete masculinization and/or fertility. Finally, it is possible that lines fully masculinized by the longer isoforms would have been recovered had we screened a much larger sample of additional individual transgenic lines.

In transgenic lines where *Nix* transgene expression of specific isoforms is strong enough, *Nix* seems to constitute a self-sufficient ectopic M-locus. Indeed, over >10 generations, the masculinizing lines remained stable, with sex-ratios not differing significantly from the wild-type strain. This result is consistent with the high flexibility of sex determination in Diptera. This flexibility is probably best illustrated in *Musca domestica* in which the Mdmd male factor is mobile between chromosomes via translocation events ^34–36^. Even though it is believed that the *Aedes* non-recombining sex-loci are currently evolving towards sex-chromosomes ^37^, this experiment shows that incipient chromosome differentiation is not essential to the species’ viability. Interestingly, the observed antero-posterior gynandromorphism after injection of a plasmid expressing CRE-recombinase into SM9 embryos shows that sex determination is tissue-autonomous in *Ae. albopictus* and is entirely consistent with the cell-autonomous function of *dsx* observed in *D. melanogaster* ^38–40^.

Finally, from an operational point of view, the masculinizing *Nix*-expressing transgenic lines could be used as genetic sexing strains, since their fluorescence genes are tightly linked to the male-determining gene. Despite SM9 pseudo-males performing well in flight assays, their mating competitiveness appeared to be 4.6-fold lower than the WT in our mating competitiveness assays. With proper adjustment in the number of males to be released in a SIT intervention ^41,42^, this line still has strong potential for field deployment. One may also be interested in measuring the competitiveness of additional, newly generated masculinizing lines since the transgene insertion may have disrupted an important gene in the lines we tested.

In conclusion, in *Aedes albopictus, Nix* is confirmed to be necessary and sufficient for complete masculinization. The developed *Nix*-expressing m/m lines carrying a reporter fluorescence transgene are a promising starting point for the development of highly stable genetic sexing strains.

## Methods

### Plasmid construction

Cloned genomic sequences were either amplified by PCR using Phusion™ High-Fidelity DNA Polymerase (ThermoFisher Scientific, France), or ordered as gBlocks from IDT DNA (Belgium). Cloning of PCR products and gBlocks in the pKSB- intermediate vector ^43^ was performed using NEBuilder® HiFi DNA Assembly (New England Biolabs, France) following the manufacturer’s instructions. Modules were then assembled by Golden Gate Cloning into a destination piggyBac vector backbone (Addgene #173496) using the Eco31I restriction enzyme (Thermo Scientific, ThermoFisher Scientific, France). Plasmids were purified using NucleoSpin Plasmid and NucleoBond Xtra Midi EF kits (Macherey Nagel, France) according to the manufacturer’s instructions. Sequence verifications were performed using Eurofins Genomics (Germany) Sanger sequencing services.

Four *Nix*-expressing piggyBac-based transgenesis plasmids were built, all made available along with their annotated nucleotide sequence at Addgene (#173505, #173665, #173666, #173667). The *Nix* endogenous promoter was captured by PCR amplification of a 2kbp region upstream of the first exon (**Supplementary Data 1**). A plasmid composed of the *Nix* endogenous promoter plus the four exons, together with an *Ae. aegypti* polyubiquitin (AePUb)-dsRed reporter gene, was built to express and track isoform 1 according to the latest terminology ^22^. A plasmid composed of the *Nix* endogenous promoter plus exons 1, 3 and 4 and an AePUb-YFP reporter gene was built to express and track isoform 2. Two plasmids encoding the *Nix* endogenous promoter, the first exon, the first intron and the second exon and the GFP reporter gene under the control of either the *OpIE2* promoter or the *AePUb* promoter were both used to express and track isoforms 3 and 4.

### Embryo microinjections

Embryo microinjections were performed as previously described for *Anopheles* eggs using a Nikon Eclipse TE2000-S inverted microscope, an Eppendorf Femtojet injector and a TransferMan NK2 micromanipulator ^43^, with injection mixes comprising 320 ng/µl of piggyBac construct and 80 ng/µl of transposase-expressing helper plasmid. We codon-optimized the sequence of piggyBac transposase for mosquitoes, added an extra nuclear localization signal to the transposase sequence, and placed it under the control of the *Aedes aegypti* PUb promoter. These changes result in a massive increase in the integration efficiency of the piggyBac construct. The plasmid has been deposited on the Addgene repository (#173404). For injections with CRE recombinase, the injected solution also contained a neutral plasmid expressing DsRed gene under the control of the PUb promoter to control for microinjection efficiency. Surviving larvae expressing eGFP and transiently showing red fluorescence were collected and examined again at the pupal stage. All larvae counting and fluorescence sorting were performed using a COPAS SELECT device (Union Biometrica, Belgium) with the provided Biosort software.

### Mosquito rearing

*Aedes albopictus* mosquitoes (strain BiA) were collected as larvae from a garden rainwater collector in the city of Bischheim, near Strasbourg (France) in 2018 and maintained in the insectary since then at 25°C, 75-80% humidity, with a 14-hr/10-hr light/dark photoperiod. Larvae were reared in pans filled with demineralized water and provided ground TetraMin fish food twice a day. Adult mosquitoes were caged and provided with 10% sugar solution *ad libitum*. Females were blood-fed on anesthetised mice. Eggs were laid three days later on wet kraft paper and allowed to develop for another three days before being dried.

### Molecular characterization of mosquito lines

Genomic DNA was extracted from pupae using NucleoSpin Tissue kit following the manufacturer’s instructions (Macherey Nagel, France). Total RNA was extracted using TRIzol RNA Isolation Reagents (Invitrogen, ThermoFisher Scientific, France) and reverse-transcribed into cDNA using RevertAid H Minus First Strand cDNA Synthesis Kit (Thermo Scientific, ThermoFisher Scientific, France). Presence of endogenous and/or exogenous *Nix* in the genome was assessed by PCR using GoTaq® Green Master Mix (M712, Promega, France) according to the manufacturer’s recommendations with the Nix-833 published primers ^25^ and primer EM1926-EM1927 (**Supplementary Table 1**). The *dsx* and *fru* splicing patterns were assessed by RT-PCR using published primers ^44^ with Phire Tissue Direct PCR Master Mix (Thermo Scientific, ThermoFisher Scientific, France). All PCRs were performed in a Veriti 96-wells Thermal Cycler (Applied Biosystems, ThermoFisher Scientific, France). Relative expression of *Nix, myo-sex*, and *myo-fem* orthologues was assessed by RT-qPCR with published primers ^22^ as well as primers provided in **Supplementary Table 1**. The AalRps7 housekeeping gene was used as a reference with published primers ^44^. qPCRs were performed using SYBR™ Fast SYBR™ Green Master Mix (ThermoFisher Scientific, France) and a 7500 Fast Real-Time PCR System machine (Applied Biosystems, ThermoFisher Scientific, France).

### Phenotypical characterization of mosquito lines

For all phenotypical tests, mosquitoes from the reference strain (BiA) and from the tested transgenic strain were hatched and reared mixed together, with the same larval density and the same amount of food in all tanks. The hatching rate was estimated by counting eggs on a piece of egg paper under a binocular microscope, submerging the eggs, placing them in a vacuum chamber for 30mn and allowing to hatch for 24h before counting the number of larvae by COPAS, on three separate egg batches. Sex ratio was measured on three samples of 100 larvae of each strain reared to pupae. Males were sorted at the pupal stage based on their fluorescence under a Nikon SMZ18 binocular microscope equipped with a Lumencor Sola Light engine. Right wings from 38 WT males, 39 WT females and 48 SM9 males were dissected under a binocular microscope and placed on a microscope slide using double-sided tape. Pictures were taken under a Zeiss SteREO binocular microscope with X-Cite Xylis engine (Excelitas technologies) and analysed using ImageJ software ^45^ to measure wing length as a proxy for body size ^27^. Male and female genitalia were observed under a binocular microscope at the pupal stage, and photographed using the Nikon SMZ18 binocular microscope. Male flight ability was assessed using a Flight Test Device as previously described, with 3 replicates for each treatment ^29^. Relative fertility was calculated by comparing the number of live larvae in the progeny from 30 males of the nix-expressing line crossed with 60 wild-type females vs. 30 wild-type males crossed with 60 wild-type females. Three independent replicates of this test were performed for each strain. Relative fecundity (total number of eggs produced by the same crosses) was estimated by dividing the total number of larvae in the progeny by the hatching rate. Relative competitiveness was estimated by placing 30 males of the nix-expressing line with 30 males of the wild-type line in a cage with 30 wild-type females in 5 independent replicates. In their progeny, we measured the percentage of *Nix*-expressing males (tracked by fluorescence using a COPAS device). If fecundity and fertility are similar in both lines, and if they were equally competitive, half the females would mate with a WT male and the other half with a transgenic male, leading to 50% WT and 50% progeny sired by transgenic males. Within the offspring of the transgenic males, the percentage of males would only depend on the line’s sex ratio. Consequently, the expected percentage of transgenic males (fluorescent larvae) in the total progeny of the competition assay would equal half the percentage of males in the transgenic strain. This theoretical value is then compared to the measured percentages of transgenic males.

### Statistical analyses

‘±’ symbols in main text represent the data standard deviation (SD) or model estimation standard errors (SE). Error bars in Figure 2 are 95% CI, while in Figure 3 they depict 5% to 95% sample distribution. The number of biological replicates (N) and type of statistical analysis are indicated in methods and figure legends. For qPCR, three technical replicates of each biological sample were used and each line was tested on three independent biological samples. To test the effects of the lines on gene expression we used linear model and normal distribution assumptions. The significance between the lines was tested by ANOVA followed by pairwise Tukey test. The effect of lines on wing length and fertility were tested using linear model and normal distribution assumptions. The effect of the lines on flight ability, hatching rates and sex ratios were tested using linear generalised mixed-effect model and binomial distribution assumptions. The replicate was set as random effect for sex ratios and flight tests as experiments were performed on different days. The relative competitiveness of transgenic lines compared to WT was tested using linear generalised mixed-effect model and binomial distribution assumptions. For this test, we compared the number of SM9 male progeny measured in the offspring of each competitiveness replicate, to the expected number of progeny that would have been obtained if both lines were as competitive. All statistical analyses and plots were performed on R software version 4.0.5 ^46^. All model assumptions were tested using the R package ‘performance’ (**Supplementary Data 3**).

## Data availability

Plasmid sequences have been deposited on Addgene (https://www.addgene.org/). All plasmids and strains described in this paper are available upon request.

## Acknowledgments

This study was funded by EU ERC CoG—682387 REVOLINC to J.B. The contents of this publication are the sole responsibility of the authors and do not necessarily reflect the views of the European Commission. Mosquito production and insectarium operation were supported by Agence Nationale de la Recherche grant #ANR-11-EQPX-0022. Part of the technical work was funded by ANR grants #ANR-19-CE35-0007 GDaMO and # 18-CE35-0003-02 BAKOUMBA to E.M.

## Conflicts of interest

The authors declare no conflict of interest.

## Notes

### Competing Interest Statement

The authors have declared no competing interest.

