## Supplementary Information for "Transgenic expression of *Nix* converts genetic females into males and allows automated sex sorting in *Aedes albopictus*"

**Supplementary Tables**

**Supplementary Table 1**: Sequences of primers used in this study.

**Supplementary Figures**

**Supplementary Figure 1:** Detection of lines composed exclusively of genetic females in lines carrying the *Nix*-OpIE2-GFP plasmid.

**Supplementary Figure 2:** Schematic of *Ae. albopictus Nix* isoforms and of their cloning in the injected plasmids.

**Supplementary Figure 3:** Intersex phenotypes in SM9 line prior to purification.

**Supplementary Figure 4:** Gallery of male SM9 pupae arising from CRE-injected embryos, showing local demasculinization in the posterior pole.

**Supplementary Figure 5:** Sex-specific *fruitless* and *doublesex* splicing patterns.

**Supplementary Data**

**Supplementary Data 1:** Sequence of the *Ae. albopictus* genomic region amplified for cloning *Nix* promoter.

**Supplementary Data 2:** Male-specific amplification pattern of one of the copies of the *myo-sex* orthologue.

**Supplementary Data 3:** Model output from all statistical analyses and performance assessment

#### Supplementary Notes

**Supplementary Note 1:** Obtaining *Nix*-expressing transgenic lines

**Supplementary Note 2:** Identification of *myo-sex* and *myo-fem* orthologues

#### References

**Supplementary Tables**

**Supplementary Table 1: Sequences of primers used in this study.**

| Name | 5' – 3' sequence | Use |
|---|---|---|
| EM1926 | CCCTCAATTTTCCGCCAACTATT | 655bp amplicon in intron 2 for detection of endogenous *Nix* |
| EM1927 | AATCTTTGGTGCGCCGTGTC | |
| Myosex369-F | AGGCCATACTAACCTTCCGT | Genomic amplification of a non-coding part of the *myo-sex* copy from scaffold 369 (end of intron 1). |
| Myosex396-R | ATACAATGAAGTAACAATGGAGCG | |
| EM2145 | CAAGTTGGTGACGATCCCGA | For RT-qPCR of mRNA from *myo-sex* orthologues |
| EM2146 | GTTGGGTAGAGCAACGGTGA | |
| EM2147 | CGCCGGAAAAACGTATCCACT | For RT-PCR of mRNA from LOC115254984, a candidate *myo-fem* orthologue |
| EM2148 | GCTGGTTCCAGGTTAGTTGG | |
| EM2149 | CCCGTGCTGAAGAGTTGGAG | For RT-PCR of mRNA from LOC115254986, a candidate *myo-fem* orthologue |
| EM2150 | GTGGACAGACGTTGCTTAGT | |
| EM2151 | GTAGGCATCTACGAGCCCAA | For RT-PCR of mRNA from LOC109402113, a candidate *myo-fem* orthologue |
| EM2152 | CCAACCTGTACCACTGGCTT | |
| EM2153 | CATTGGAAACATTCCCGCCG | For RT-PCR or RT-qPCR of mRNA from *Nix* |
| EM2154 | ACTGCCGGTTTCACATCACA | |
| EM2170 | ACGTGCCGAAGAATTGGAAG | For RT-qPCR of mRNA from LOC109402113, putative *myo-fem* orthologue |
| EM2171 | TTCTAAGGCAACACACTTCTGA | |
| EM2174 | CGTGCCACCCTTCTTGGTAA | For RT-qPCR of mRNA from LOC115254984, candidate *myo-fem* orthologue |
| EM2175 | CCTCCAACTCTTCTGCACGG | |

**Supplementary Figures**

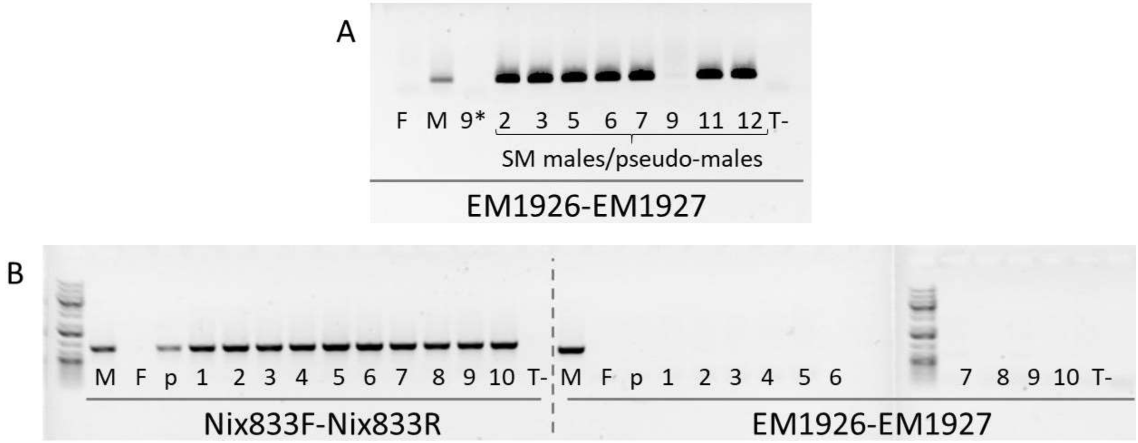


**Supplementary Figure 1: Detection of lines composed exclusively of genetic females in lines carrying the *Nix*-OpIE2-GFP plasmid.**

**A)** PCR amplification of genomic DNA from pooled phenotypic males using primers EM1926-EM1927. This primer pair is located within the second intron of *Nix*, absent from our construct, and thus amplifies only DNA from genetic males. Genomic DNA was extracted from pooled pupae and analysed in the following order: WT females ("F", negative control), WT males ("M", positive control), SM9 partially masculinized female ("9*"), males or pseudo-males from selected lines arising from injection of the first OpIE2-GFP -marked *Nix* expressing plasmid (numbers 2 to 12), negative control without DNA template ("T-"). This PCR allowed identification of line SM9 as composed exclusively of genetic females. **B)** Screening of 10 SM9 individual males (labelled 1 to 10) with primer pair Nix-833F-Nix-833R amplifying any *Nix*-bearing genomic DNA [1] and primer pair EM1926-EM1927 for endogenous *Nix* only. The following controls were used: "M" is WT male positive control, "F" is WT female negative control, "p" is the pooled SM9 male DNA from top panel, and "T-" is a negative control without template DNA. This PCR confirmed that all phenotypic males in the SM9 line were pseudo-males. PCR products were resolved on a 0.8% agarose gel with 0.2µg/mL ethidium bromide.

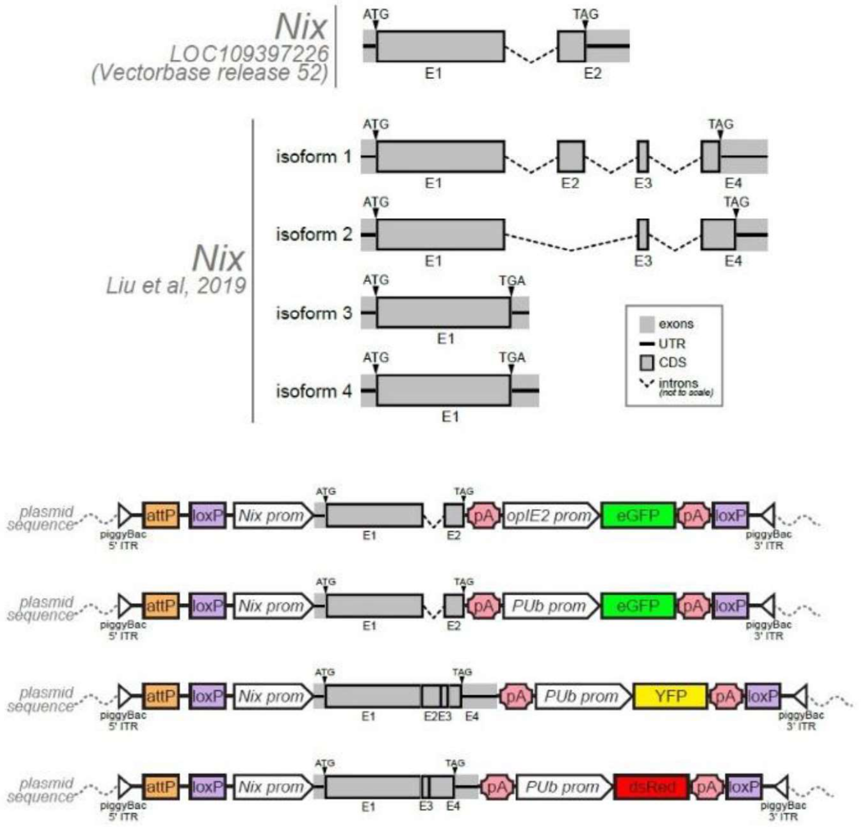


**Supplementary Figure 2: Schematic of *Ae. albopictus Nix* isoforms and of their cloning in the injected plasmids.**

The top part represents previous [1] and current [2] knowledge of the *Ae. albopictus* Nix gene structure. Isoform naming follows work published in [2]. Grey boxes are *Nix* exons. The bottom part schematizes the plasmids built in this work. White triangles are piggyBac 5' and 3' inverted terminal repeat (ITR) sequences, orange boxes represent an AttP landing site included for potential future purposes, purple boxes represent loxP recombination sites, white arrows are promoters, grey boxes are *Nix* exons, pink polygons are SV40 polyA sequences, green, yellow and red boxes are eGFP, YFP and DsRed gene sequences, respectively. Drawing not to scale. Detailed sequences can be found under Addgene references #173505, #173665, #173666, #173667.

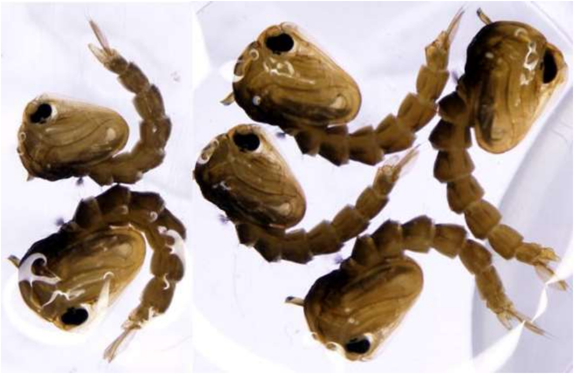

**Supplementary Figure 3: Intersex phenotypes in SM9 line prior to purification.**

The two pupae on the left side of the picture are a control male and a control female from SM9 line. All four pupae on the right are representative GFP-expressing intersex individuals with deformed genitalia found in the SM9 line prior to elimination of additional non-fully masculinizing transgene insertions. Pictures were taken under a binocular microscope with white light.

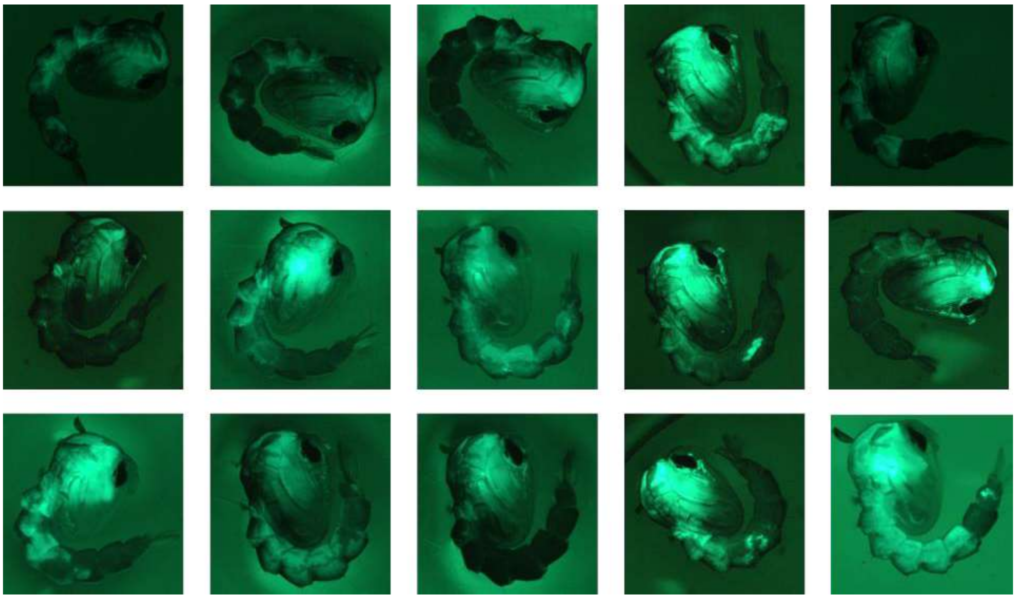

**Supplementary Figure 4: Gallery of male SM9 pupae arising from CRE-injected embryos, showing local demasculinization in the posterior pole.**

All adults that emerged from these pupae had male heads and female genitalia. Pictures were taken under a binocular fluorescence microscope using a GFP filter.

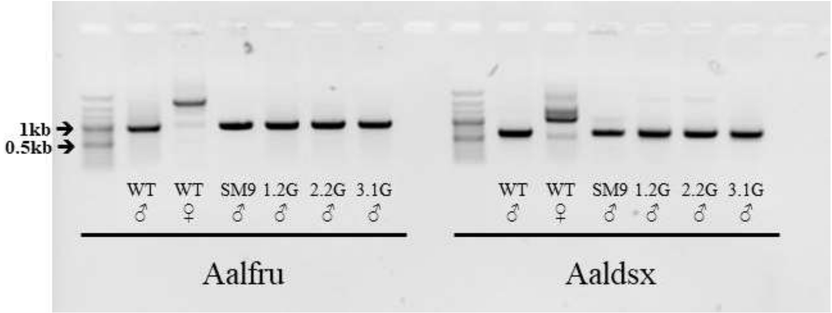


**Supplementary Figure 5: Sex-specific *fruitless* and *doublesex* splicing patterns.**

mRNAs from WT male, WT female, SM9 pseudo-male, 1.2G pseudo-male, 2.2G pseudo-male, and 3.1G pseudo-male pupae were extracted, reverse-transcribed into cDNA, which was used for PCR. Products were resolved on a 1.5% agarose gel with 0.2µg/mL ethidium bromide. Left side: RT-PCR with *fruitless* sex-specific primers [3] for which the male product is expected at 987 bp and the female product at 2010 bp. Right side: RT-PCR with *doublesex* primers [3], for which the male product is expected at 620 bp and the female product at 1062 bp.

**Supplementary Data**

**Supplementary Data 1: Sequence of the *Ae. albopictus* genomic region amplified for cloning *Nix* promoter.**

In the sequence below, *Nix* promoter sequence is included in the 1,916 bp sequence highlighted in grey, the 5' UTR sequence of the *Nix* gene is not highlighted, while the ATG start codon is highlighted in green.

TCGCATTTTATGAGTAAAGGCCCATTACTCATATATGGGTAAAGTGCTTTTTGTAAAAAAT

GAGTAAATCGATTTACTCATAAATTATAATTTCACCGTGTTTACTTTTCGTCTTTTGTTATA

TTTTTGAACTGTTAACTATTTTGTATTTATCATTTAATTCATACAAAATTTTGCGTTTATTT

TGTAGTTTATTGCAATCTACGAAAATTATTAATGAATTTTTACATGTTTCAAATACTCTTA

ATCATTTTCCTCCTGGAAAACAGACCTTCCGAGACGTAGCTCTTTGAAGTTTTCATAATGA

TCCATGTATGAAATTTTCCCCTCCGGCTGTTGTTCATCGCGCAGTTGTTCCTGTTCTTCCGT

TCCTCGTTCTTCCGGCGCTTGAAGCCTTGGTGCAGTCGGAACTGTGGAAAAGACTGATAT

ATTTTTATTTCACGGTCGTTAATGAAAATGATGCTGTCACTATGGAACTTACCTGCAAATT

CCATGCTTTTGTAGCCTTTCTTGTAGCTCCACATGATGATTTGTTGCTCCTGTATTATCTTC

GAGTACCAAACCTTGTGAAACGGAAATAATCCATCGTAAATGAAAATTGAACGGTATCA

ATTTATCCAGGAAGTACTTACCGATTAAAATATCAATGTATTTGGGAACCTCGACTCTAAT

CCGCGTGGGAATCCGATGGACGCGGTATCGGTCATTGAACCCGATGCTGAAGGGCCTCGA

CTGGCTATAGCGGTGCCGTGAAAATCAAAATATCTAAAATTTCCCGATAGTGTTATGATC

AGGGGATCCGCATGCGGCAAGGTAGGCCTACTGCTCATCTTTTGCGAGTGTGTTAGTGTG

ACTGGCAACGCGACCAGAAGTCTAGTCGAATCCTGCTTGGAAATCGGTTTGTTTTGTCGA

TGGGGGTGGCCCGCGAAATCGCATTGTGTTGTTTGGGAAGGCCCGCTCGATGCATGTTAG

GTAGGCGTAGTTTGTTCGAGTTTGACGTGCCTGCACCCTCCCTGGTTATGATGGCGCACTA

ATGCATTCCGAGAAAAGTTGCTAACTTGAAAATTTTCATCCAAATATGGGTAAGTTCATTT

ACCCATGTTTGGGTATAGCGATTTTACCTATAATATGGGCAAATTTGACCGTTTTTCAAAG

GTATATTTTACCCATATTATGGGTAATGCATATGAAACCCAAATATGGGTAAATCAACTA

CTGATCTATGGGTAAACTGATCTTAGCGTGTATGTTTCACACGTGACATTCACTGGTTTTA

CAATTGGCTTCTTTAGCAGTGTTGGGATAATTCCATATTATGAATCTGCATGCCAAACTGG

GCCGAAATCCAAATTTTCATCAATTTTGGTGCACGGGAACCTATTTAAATATCAATTTGAA

GTTTGTATGGGAGCGATTTGTCGAATCACCCCTCGTTGCATTTTGTACTGGGCGGAGCTGT

CAAACAGTTGCCTAGCTGTCAAAAGGTGATTTCGAATAATCTCTTTGAAATTGATTTTAGG

TATCAAAATAAAGTTCTAAAAATCTGAAAAAAATCATAGTGGCTCAGAAAAAGGTGCTCT

TTCGTATAAAATCAAAAAATGAACACTTTTTTCAAAATTTAAAAACCCAATTTTCGCAAA

TCTATAGGGCTCTGCACGAAGTTCTCTCCCTCTCTTTCGCTCTCATTGAGATTTTGTAAAC

AACAAGGCCAGGAAATGTCAAAATCCCATACAAAATCAAAACAGTGCAGTGCCCTATAT

GTAAAACACATCACTCCGACGTGTAAATTTTTTTGAGTGTTGATTTAATCAAAGTGAATA

AAAATATTAGTTTTATGACATACTTGTTTTCTGAGTGTAGCAAAAATATGAAAACACATTT

TTGTACTTTGAATGTTAAGCGTGTATGCTTTTTGTGTCAATATGTCAATTGTTAAACCCAT

GTTAATAGTTTTAATTTTTTTTTAATCAAATTCTTTTTTTAAGTAATG

**Supplementary Data 2: Male-specific amplification pattern of one of the copies of the *myo-sex* orthologue.**

PCR amplification was performed using primers Myosex369-F and Myosex369-R targeting the end of the first intron of *myo-sex*. The expected product with this primer pair was 997 bp as provided below. The discovered male-specific product carries a 664 bp deletion (highlighted in grey) and is present in genetic males only.

AGGCCATACTAACCTTCCGTAAATCACGCGGTTGGCCACTACCGTCAGCACCACGATAAT
GCTGAGTACAAAATTTAGATTTCTTCAACGTGTTGTACAAAATACAAAAGCGCGATCGCT
CGAGGGCTAAGAATTAATGAAGAAAATTGCAAATCAATTACATATTTATACAAAATCATA
ACAATAGGTATAGAT<u>G</u>TTTGCAAAGATGTTCGTTGACAGCTTGAATCTCGAAACAGAGAT
GAATAGATTGAACTATAAAACAAACTCGATCAAAAATAACACAGCGTAACAAAAATAAC
TTTTTGTATGTCTCTAGAGCAAACTTACGTGTCTCCGAAGGATTTTGGGCCGCTGAATCCG
AATCTGGGCTCAGATTTGCTCTAACACGTCACAATTTTGAGCTATACCTCAATTTATAGGG
CAAAATATGCGATTTTGGGCTTTTTTGACTGCAAGCCATTAAGCAAGGAAATATTTTTTTA
AGCAATCAAAAGGTTAATTGGTCAATTAACATCTAAATTACGACTCATGCAAAATATTTC
GTTTTACCAAATCGAATTTGATAGTTTTAAGCGATTTATGTTAGGTACGATATTTCCCATA
TAAGTCAGCCTCCAAAAGTTGCATGCAAGTTTTCATGCTAACATAAAATGCTTAAATCCA
TCAAATTTGATTAGGTAAAACGAAATATTTTGCATGAGTCGTTAATTTAGATGTTAGTTGA
CCAATTAACCTTTTGATTGCTTAAAAAAAATATTTCCTTGCTTAATGGCTTGCAGTCAAAA
AAGCCCAAAATCGCATATTTTGCCCTATAAATTGAGGTATAGCTCAAAATTGTGACGTGT
TAGAGCAAATCTAAGCCCGGATTCGGATTCAGCGGCCCAAAATCCTTCGGAGACACATAA
GTTTGCTCTTGAGACAAAAAAAATGTTGCGCTGTGTAATCTGTTAAAAACATTGTCCTAC
ATTTTTTGCGCTCCATTGTTACTTCATTGTAT

**Supplementary Data 3: Model output from all statistical analyses and performance assessment**

### Supplementary material : model outputs

#### Sex ratios

##### Goal

To evaluate whether the Nix-eGFP cassette was stable and adult males fully viable, we first determined the sex ratios of transgenic strains in comparison to that of the parental wild strain (WT).

##### Method

The effect of the lines on sex ratios was tested using linear generalised mixed-effect model and binomial distribution assumptions. The replicate was set as random effect as mesurements were performed on different days. The significance between the lines was tested by ANOVA followed by pairwise Tukey test.

##### Model output

```
## Generalized linear mixed model fit by maximum likelihood (Laplace
##   Approximation) [glmerMod]
##  Family: binomial  ( logit )
## Formula: male_percentage ~ line + (1 | replicate)
##    Data: males
##
##      AIC      BIC   logLik deviance df.resid
##  21048.9  21087.1 -10519.5  21038.9    15226
##
## Scaled residuals:
##     Min      1Q  Median      3Q     Max
## -1.1670 -1.0378  0.8569  0.9635  1.0616
##
## Random effects:
##  Groups    Name        Variance Std.Dev.
##  replicate (Intercept) 0.004062 0.06374
## Number of obs: 15231, groups:  replicate, 3
##
## Fixed effects:
##             Estimate Std. Error z value Pr(>|z|)
## (Intercept) -0.05054    0.04544  -1.112    0.266
## line3.1G     0.19383    0.03978   4.873 1.10e-06 ***
## lineBiA      0.12214    0.09563   1.277    0.202
## lineSM9      0.28682    0.04367   6.568 5.09e-11 ***
## ---
## Signif. codes:  0 '***' 0.001 '**' 0.01 '*' 0.05 '.' 0.1 ' ' 1
##
## Correlation of Fixed Effects:
##          (Intr) ln3.1G lineBA
## line3.1G -0.377
## lineBiA  -0.189  0.205
## lineSM9  -0.376  0.422  0.219
```

```
##
##   Simultaneous Tests for General Linear Hypotheses
##
## Multiple Comparisons of Means: Tukey Contrasts
##
##
## Fit: glmer(formula = male_percentage ~ line + (1 | replicate), data = males,
##     family = binomial(link = logit))
##
## Linear Hypotheses:
##                  Estimate Std. Error z value Pr(>|z|)
## 3.1G - 1.2G == 0  0.19383    0.03978   4.873   <0.001 ***
## BiA - 1.2G == 0   0.12214    0.09563   1.277    0.558
## SM9 - 1.2G == 0   0.28682    0.04367   6.568   <0.001 ***
## BiA - 3.1G == 0  -0.07169    0.09573  -0.749    0.869
## SM9 - 3.1G == 0   0.09299    0.04500   2.067    0.151
## SM9 - BiA == 0    0.16468    0.09604   1.715    0.297
## ---
## Signif. codes:  0 '***' 0.001 '**' 0.01 '*' 0.05 '.' 0.1 ' ' 1
## (Adjusted p values reported -- single-step method)
```

```
##   Line  Estimate  Std_Error
## 1 1.2G 0.4873675 0.04544079
## 2 3.1G 0.5357612 0.04779783
## 3  BiA 0.5178926 0.09781716
## 4  SM9 0.5587964 0.04981261
```

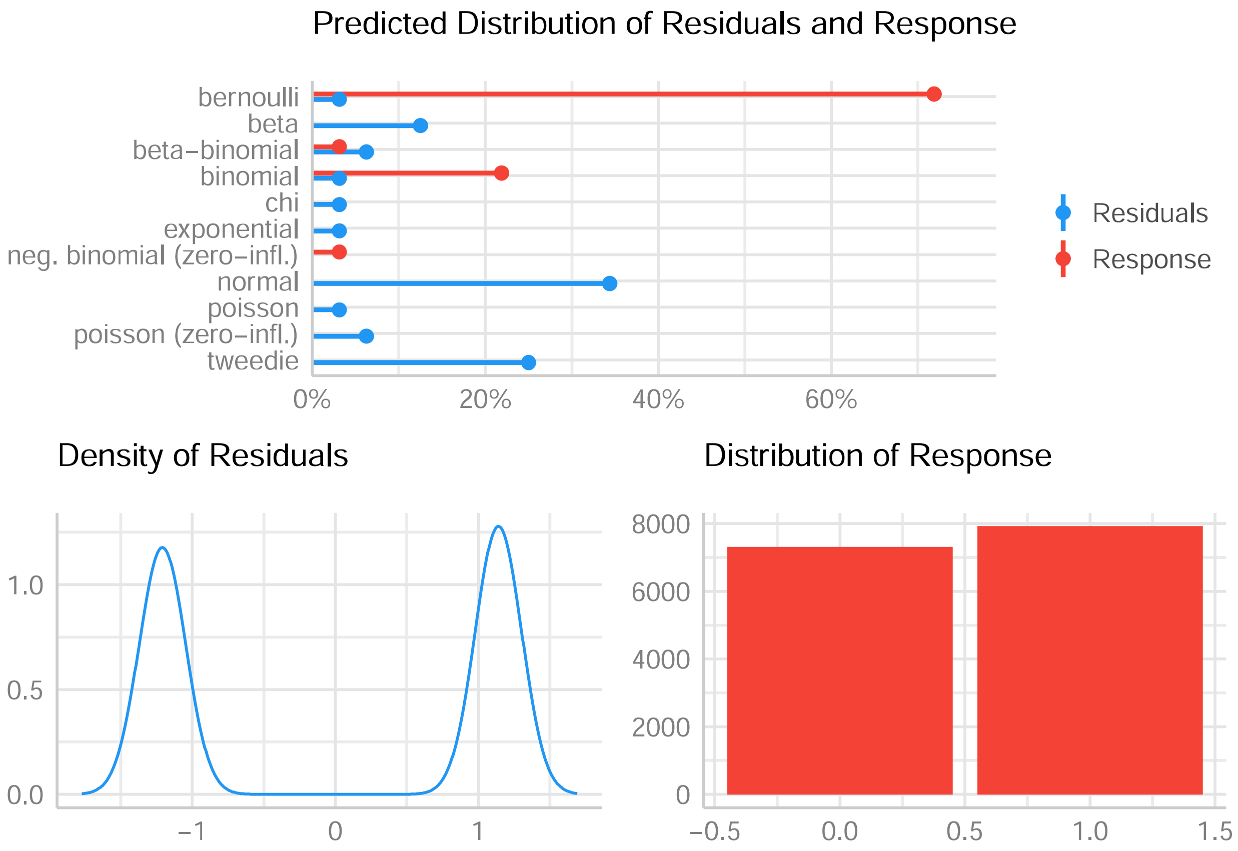

### Wing lengths

#### Goal

In *Aedes* mosquitoes, males and females display a significant size dimorphism, with females having a larger body size. We compared the body size of masculinized genetic females to WT females and WT males using wing length as a proxy for mosquito body size.

#### Method

The effect of lines on wing length was tested using linear model and normal distribution assumptions. The significance between the lines was tested by ANOVA followed by pairwise Tukey test.

#### Model output

```
##
## Call:
## lm(formula = Length ~ Line, data = wings)
##
## Residuals:
##     Min      1Q  Median      3Q     Max
## -71.257 -15.393  -1.055  16.187 127.278
##
## Coefficients:
##              Estimate Std. Error t value Pr(>|t|)
## (Intercept)   815.065      4.469  182.40   <2e-16 ***
## LineBiA male -109.314      6.361  -17.18   <2e-16 ***
## LineSM9 male -108.970      6.016  -18.11   <2e-16 ***
## ---
## Signif. codes:  0 '***' 0.001 '**' 0.01 '*' 0.05 '.' 0.1 ' ' 1
##
## Residual standard error: 27.91 on 122 degrees of freedom
## Multiple R-squared:  0.7708, Adjusted R-squared:  0.767
## F-statistic: 205.1 on 2 and 122 DF,  p-value: < 2.2e-16
```

```
##   Tukey multiple comparisons of means
##     95% family-wise confidence level
##
## Fit: aov(formula = mod.wings)
##
## $Line
##                              diff        lwr       upr     p adj
## BiA male-BiA female -109.3141235 -124.40642 -94.22182 0.0000000
## SM9 male-BiA female -108.9702244 -123.24407 -94.69638 0.0000000
## SM9 male-BiA male      0.3438991  -14.03319  14.72099 0.9982258
```

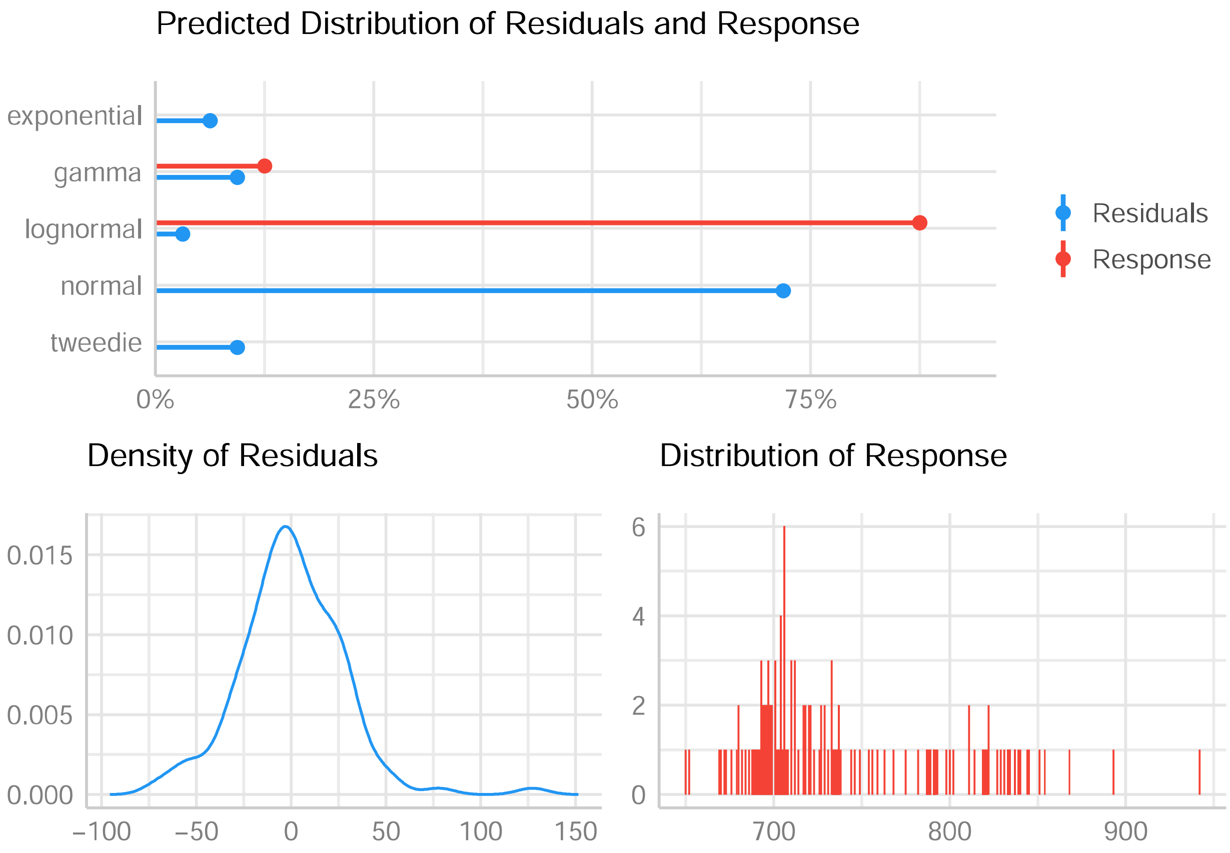

Predicted Distribution of Residuals and Response
exponential
gamma
lognormal
normal
tweedie
0%
25%
50%
75%
Residuals
Response
Density of Residuals
0.015
0.010
0.005
0.000
−100
−50
0
50
100
150
Distribution of Response
6
4
2
0
700
800
900

### RT-qPCR : relative expresion of *Nix*

#### Goal

To assess whether transgenic males express *Nix* in similar levels to wild-type males, expression levels were measured by RT-qPCR at the pupal stage.

#### Method

For qPCR, three technical replicates of each biological sample were used. Each line was tested on 3 independent biological samples. To test the effects of the lines on gene expression we used linear model and normal distribution assumptions.The significance between the lines was tested by ANOVA followed by pairwise Tukey test.

#### Model output

```
##
## Call:
## lm(formula = dCt ~ line, data = df.nix)
##
## Residuals:
##     Min      1Q  Median      3Q     Max
## -2.7189 -1.2011  0.1234  0.9289  2.7189
##
## Coefficients:
##               Estimate Std. Error t value Pr(>|t|)
## (Intercept)     -3.957      1.116  -3.547  0.00625 **
## lineWT female  -10.653      1.578  -6.753 8.34e-05 ***
## lineSM9 male    -2.923      1.764  -1.657  0.13190
## line1.2G male   -2.964      1.578  -1.879  0.09298 .
## line3.1G male    2.255      1.578   1.429  0.18676
## ---
## Signif. codes:  0 '***' 0.001 '**' 0.01 '*' 0.05 '.' 0.1 ' ' 1
##
## Residual standard error: 1.932 on 9 degrees of freedom
##   (4 observations deleted due to missingness)
## Multiple R-squared:  0.8946, Adjusted R-squared:  0.8478
## F-statistic:  19.1 on 4 and 9 DF,  p-value: 0.0002012
```

```
##   Tukey multiple comparisons of means
##     95% family-wise confidence level
##
## Fit: aov(formula = mod.dCt)
##
## $line
##                             diff         lwr       upr     p adj
## WT female-WT male   -10.65328068 -15.9581933 -5.348368 0.0005953
## SM9 male-WT male     -2.92265574  -8.8537284  3.008417 0.5014045
## 1.2G male-WT male    -2.96413962  -8.2690522  2.340773 0.3910419
## 3.1G male-WT male     2.25456640  -3.0503462  7.559479 0.6262636
## SM9 male-WT female    7.73062494   1.7995523 13.661698 0.0115720
## 1.2G male-WT female   7.68914106   2.3842284 12.994054 0.0059238
## 3.1G male-WT female  12.90784709   7.6029345 18.212760 0.0001348
## 1.2G male-SM9 male   -0.04148388  -5.9725565  5.889589 0.9999999
## 3.1G male-SM9 male    5.17722214  -0.7538505 11.108295 0.0933632
```

```
## 3.1G male-1.2G male   5.21870602  -0.0862066 10.523619 0.0541624
```

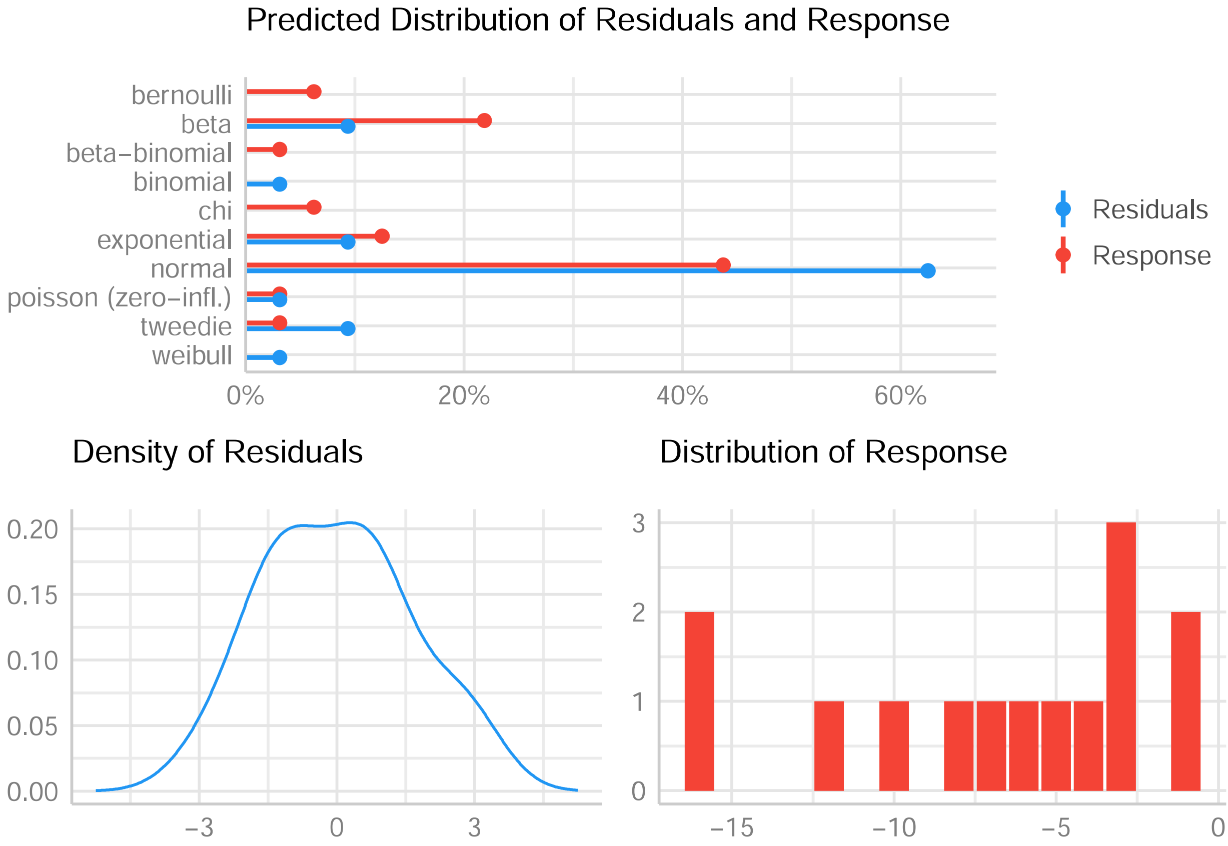

### RT-qPCR : relative expresion of *myo-sex*

#### Goal

To assess whether transgenic males express *myo-sex* in similar levels to wild-type males, expression levels were measured by RT-qPCR at the pupal stage.

#### Method

For qPCR, three technical replicates of each biological sample were used. Each line was tested on 3 independent biological samples. To test the effects of the lines on gene expression we used linear model and normal distribution assumptions.The significance between the lines was tested by ANOVA followed by pairwise Tukey test.

#### Model output

```
##
## Call:
## lm(formula = dCt ~ line, data = df.myosex)
##
## Residuals:
##      Min       1Q   Median       3Q      Max
## -1.33657 -0.44239  0.06005  0.37782  1.13212
##
## Coefficients:
##               Estimate Std. Error t value Pr(>|t|)
## (Intercept)    3.69459    0.49719   7.431 3.97e-05 ***
## lineWT female -4.50142    0.70313  -6.402 0.000125 ***
## lineSM9 male   0.07876    0.78613   0.100 0.922396
## line1.2G male  0.46635    0.70313   0.663 0.523796
## line3.1G male  0.71218    0.70313   1.013 0.337575
## ---
## Signif. codes:  0 '***' 0.001 '**' 0.01 '*' 0.05 '.' 0.1 ' ' 1
##
## Residual standard error: 0.8612 on 9 degrees of freedom
## Multiple R-squared:  0.8937, Adjusted R-squared:  0.8464
## F-statistic: 18.91 on 4 and 9 DF,  p-value: 0.0002093
```

```
##   Tukey multiple comparisons of means
##     95% family-wise confidence level
##
## Fit: aov(formula = mod.dCt)
##
## $line
##                           diff       lwr       upr     p adj
## WT female-WT male   -4.50142023 -6.865756 -2.137084 0.0008858
## SM9 male-WT male     0.07875527 -2.564652  2.722163 0.9999723
## 1.2G male-WT male    0.46634928 -1.897986  2.830685 0.9596094
## 3.1G male-WT male    0.71218332 -1.652152  3.076519 0.8434409
## SM9 male-WT female   4.58017550  1.936768  7.223583 0.0017537
## 1.2G male-WT female  4.96776952  2.603434  7.332105 0.0004226
## 3.1G male-WT female  5.21360355  2.849268  7.577939 0.0002916
## 1.2G male-SM9 male   0.38759401 -2.255814  3.031002 0.9860491
## 3.1G male-SM9 male   0.63342805 -2.009980  3.276836 0.9223992
## 3.1G male-1.2G male  0.24583403 -2.118502  2.610170 0.9961853
```

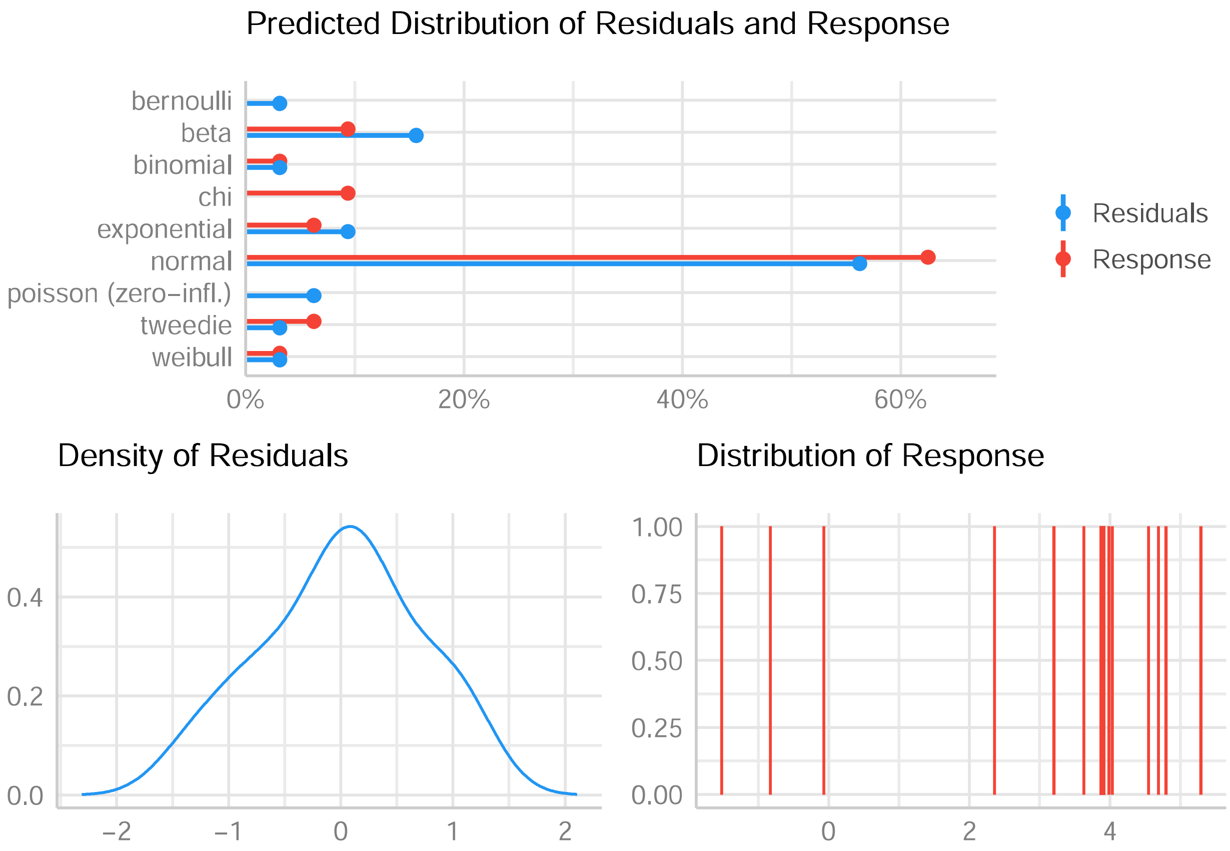
Predicted Distribution of Residuals and Response
bernoulli
beta
binomial
chi
exponential
normal
poisson (zero−infl.)
tweedie
weibull
0%
20%
40%
60%
Residuals
Response
Density of Residuals
0.0
0.2
0.4
−2
−1
0
1
2
Distribution of Response
0.00
0.25
0.50
0.75
1.00
0
2
4

### RT-qPCR : relative expresion of LOC109402113

#### Goal

To assess whether transgenic males express LOC109402113 (*Ae. aegypti myo-fem* homologue) in similar levels to wild-type males, expression levels were measured by RT-qPCR at the pupal stage.

#### Method

For qPCR, three technical replicates of each biological sample were used. Each line was tested on 3 independent biological samples. To test the effects of the lines on gene expression we used linear model and normal distribution assumptions.The significance between the lines was tested by ANOVA followed by pairwise Tukey test.

#### Model output

```
##
## Call:
## lm(formula = dCt ~ line, data = df.myofem70)
##
## Residuals:
##     Min      1Q  Median      3Q     Max
## -1.4024 -0.5682 -0.1063  0.2026  2.1966
##
## Coefficients:
##               Estimate Std. Error t value Pr(>|t|)
## (Intercept)   -12.2530     0.7253 -16.894 4.00e-08 ***
## lineWT female  12.4843     1.0257  12.171 6.82e-07 ***
## lineSM9 male    0.4245     1.1468   0.370    0.720
## line1.2G male   1.0439     1.0257   1.018    0.335
## line3.1G male   1.4883     1.0257   1.451    0.181
## ---
## Signif. codes:  0 '***' 0.001 '**' 0.01 '*' 0.05 '.' 0.1 ' ' 1
##
## Residual standard error: 1.256 on 9 degrees of freedom
## Multiple R-squared:  0.9584, Adjusted R-squared:  0.9399
## F-statistic: 51.86 on 4 and 9 DF,  p-value: 3.238e-06
```

```
##   Tukey multiple comparisons of means
##     95% family-wise confidence level
##
## Fit: aov(formula = mod.dCt)
##
## $line
##                            diff        lwr       upr     p adj
## WT female-WT male    12.4843140   9.035306 15.933322 0.0000051
## SM9 male-WT male      0.4245338  -3.431574  4.280642 0.9952514
## 1.2G male-WT male     1.0438734  -2.405134  4.492881 0.8412790
## 3.1G male-WT male     1.4883477  -1.960660  4.937356 0.6139971
## SM9 male-WT female  -12.0597801 -15.915888 -8.203672 0.0000174
## 1.2G male-WT female -11.4404406 -14.889448 -7.991433 0.0000106
## 3.1G male-WT female -10.9959663 -14.444974 -7.546958 0.0000148
## 1.2G male-SM9 male    0.6193395  -3.236768  4.475448 0.9805259
## 3.1G male-SM9 male    1.0638138  -2.792294  4.919922 0.8792667
## 3.1G male-1.2G male   0.4444743  -3.004534  3.893482 0.9913742
```

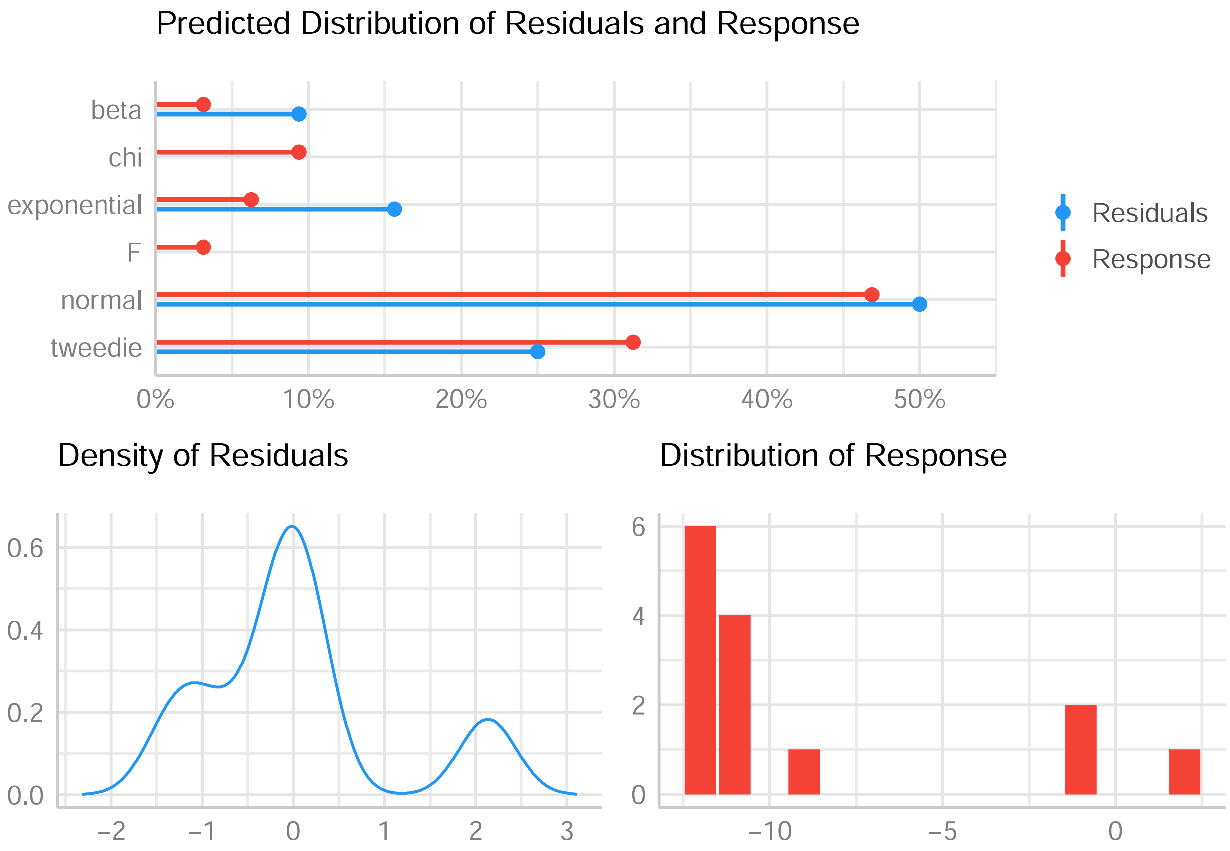

Predicted Distribution of Residuals and Response
beta
chi
exponential
F
normal
tweedie
0%
10%
20%
30%
40%
50%
Residuals
Response
Density of Residuals
0.6
0.4
0.2
0.0
−2
−1
0
1
2
3
Distribution of Response
6
4
2
0
−10
−5
0

### RT-qPCR : relative expresion of LOC115254984

#### Goal

To assess whether transgenic males express LOC115254984 (*Ae. aegypti myo-fem homologue*) in similar levels to wild-type males, expression levels were measured by RT-qPCR at the pupal stage.

#### Method

For qPCR, three technical replicates of each biological sample were used. Each line was tested on 3 independent biological samples. To test the effects of the lines on gene expression we used linear model and normal distribution assumptions.The significance between the lines was tested by ANOVA followed by pairwise Tukey test.

#### Model output

```
##
## Call:
## lm(formula = dCt ~ line, data = df.myofem74)
##
## Residuals:
##     Min      1Q  Median      3Q     Max
## -2.1395 -0.9368 -0.5869  0.6792  3.5800
##
## Coefficients:
##               Estimate Std. Error t value Pr(>|t|)
## (Intercept)   -14.9030     1.1233 -13.267 9.94e-07 ***
## lineWT female  12.6737     1.5886   7.978 4.46e-05 ***
## lineSM9 male   -1.1926     1.7762  -0.671    0.521
## line1.2G male   2.1107     1.7762   1.188    0.269
## line3.1G male   0.6873     1.5886   0.433    0.677
## ---
## Signif. codes:  0 '***' 0.001 '**' 0.01 '*' 0.05 '.' 0.1 ' ' 1
##
## Residual standard error: 1.946 on 8 degrees of freedom
## Multiple R-squared:  0.9224, Adjusted R-squared:  0.8836
## F-statistic: 23.77 on 4 and 8 DF,  p-value: 0.0001703
```

```
##   Tukey multiple comparisons of means
##     95% family-wise confidence level
##
## Fit: aov(formula = mod.dCt)
##
## $line
##                            diff        lwr       upr     p adj
## WT female-WT male    12.6736647   7.185298 18.162031 0.0003073
## SM9 male-WT male     -1.1926133  -7.328794  4.943567 0.9573246
## 1.2G male-WT male     2.1106772  -4.025503  8.246858 0.7580825
## 3.1G male-WT male     0.6872514  -4.801115  6.175618 0.9912738
## SM9 male-WT female  -13.8662780 -20.002459 -7.730097 0.0003583
## 1.2G male-WT female -10.5629875 -16.699168 -4.426807 0.0022847
## 3.1G male-WT female -11.9864133 -17.474780 -6.498047 0.0004555
## 1.2G male-SM9 male    3.3032905  -3.418558 10.025140 0.4846920
## 3.1G male-SM9 male    1.8798647  -4.256316  8.016045 0.8221682
## 3.1G male-1.2G male  -1.4234258  -7.559606  4.712755 0.9230466
```

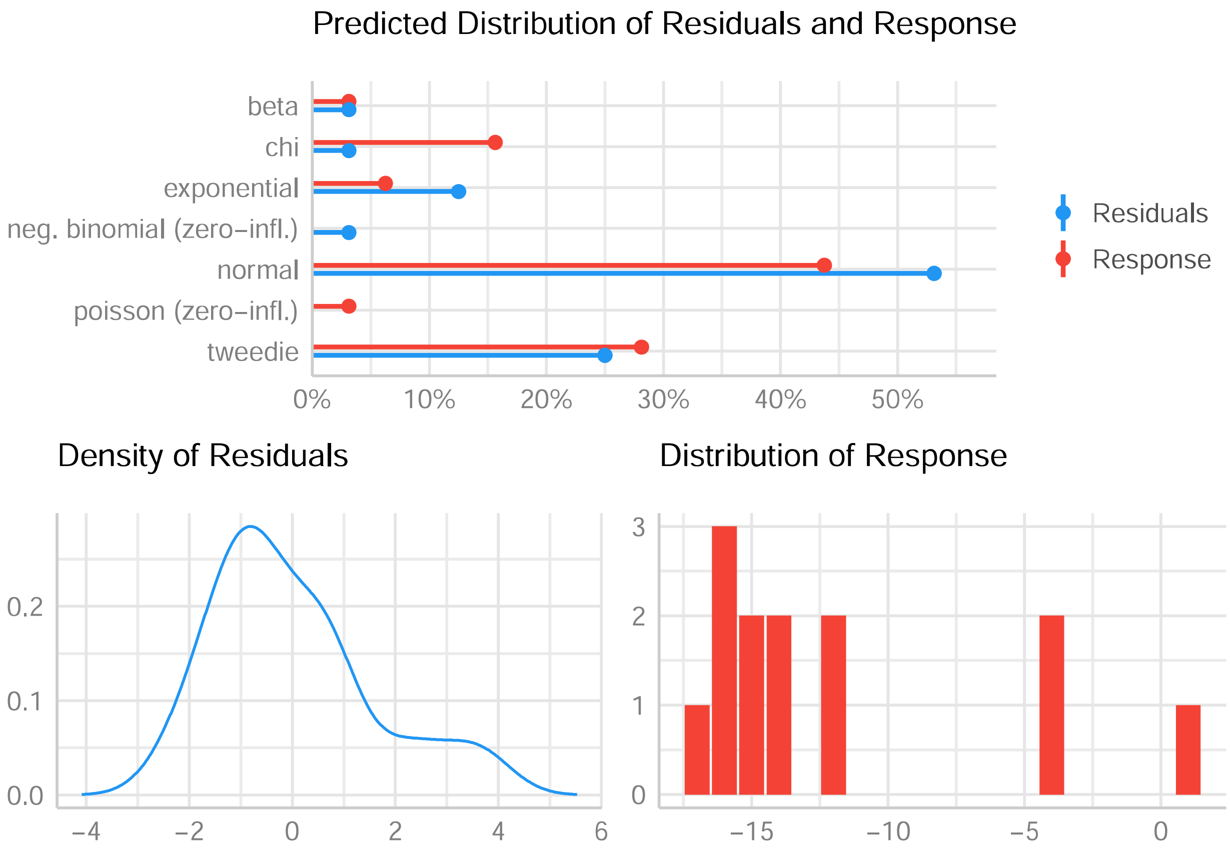
Predicted Distribution of Residuals and Response
beta
chi
exponential
neg. binomial (zero−infl.)
normal
poisson (zero−infl.)
tweedie
0%
10%
20%
30%
40%
50%
Residuals
Response
Density of Residuals
0.2
0.1
0.0
−4
−2
0
2
4
6
Distribution of Response
3
2
1
0
−15
−10
−5
0

### Flight ability

#### Goal

Since we observed that genes potentially involved in flight were regulated similarly in *Nix*-expressing pseudo-males comparing to wild-type counterparts, we compared SM9 males' flight ability to WT males by performing a flight test.

#### Method

The effect of the lines on flight ability was tested using linear generalized mixed-effect model and binomial distribution assumptions. The replicate was set as random effect for flight tests as experiments was performed on different days.

#### Model output

```
## Generalized linear mixed model fit by maximum likelihood (Laplace
##   Approximation) [glmerMod]
##  Family: binomial  ( logit )
## Formula: escape.rate ~ line + (1 | replicate)
##    Data: flight
##
##      AIC      BIC   logLik deviance df.resid
##    599.5    612.0   -296.7    593.5      486
##
## Scaled residuals:
##     Min      1Q  Median      3Q     Max
## -1.3666 -0.7490 -0.3862  0.9328  2.5890
##
## Random effects:
##  Groups    Name        Variance Std.Dev.
##  replicate (Intercept) 0.625    0.7906
## Number of obs: 489, groups:  replicate, 3
##
## Fixed effects:
##             Estimate Std. Error z value Pr(>|z|)
## (Intercept)  -0.8617     0.4796  -1.797 0.072351 .
## lineSM9       0.7171     0.1996   3.593 0.000327 ***
## ---
## Signif. codes:  0 '***' 0.001 '**' 0.01 '*' 0.05 '.' 0.1 ' ' 1
##
## Correlation of Fixed Effects:
##         (Intr)
## lineSM9 -0.217
```

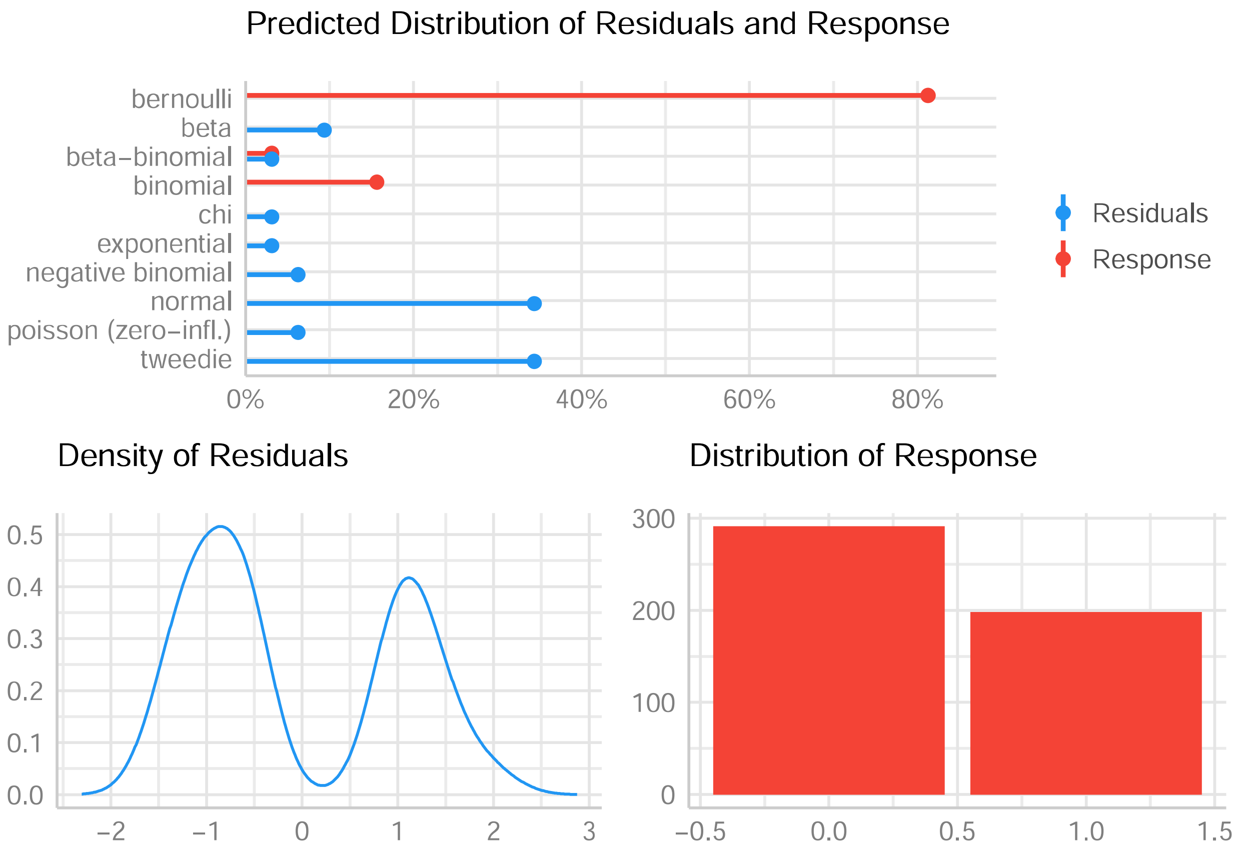
Predicted Distribution of Residuals and Response
bernoulli
beta
beta−binomial
binomial
chi
exponential
negative binomial
normal
poisson (zero−infl.)
tweedie
0%
20%
40%
60%
80%
Residuals
Response
Density of Residuals
0.5
0.4
0.3
0.2
0.1
0.0
−2
−1
0
1
2
3
Distribution of Response
300
200
100
0
−0.5
0.0
0.5
1.0
1.5

### Hatching rate

#### Goal

We compared the hatching rate of eggs produced by the trasngenic SM9 line and the control wild-type line.

#### Method

The effect of the lines on hatching rates was tested using linear generalized mixed-effect model and binomial distribution assumptions.

#### Model output

```
##
## Call:
## glm(formula = "hatched ~ line", family = binomial(link = logit),
##     data = hatch)
##
## Deviance Residuals:
##    Min      1Q  Median      3Q     Max
## -1.271  -1.271   1.086   1.086   1.118
##
## Coefficients:
##             Estimate Std. Error z value Pr(>|z|)
## (Intercept)  0.21806    0.05930   3.677 0.000236 ***
## lineSM9     -0.07723    0.09635  -0.802 0.422842
## ---
## Signif. codes:  0 '***' 0.001 '**' 0.01 '*' 0.05 '.' 0.1 ' ' 1
##
## (Dispersion parameter for binomial family taken to be 1)
##
##     Null deviance: 2545.5  on 1847  degrees of freedom
## Residual deviance: 2544.8  on 1846  degrees of freedom
## AIC: 2548.8
##
## Number of Fisher Scoring iterations: 3
```

```
## # A tibble: 2 x 3
##   line  estimate     SE
##   <chr>    <dbl>  <dbl>
## 1 BiA      0.554 0.0147
## 2 SM9      0.535 0.0189
```

```
## Warning: Removed 9 rows containing missing values (geom_segment).
```

```
## Warning: Removed 9 rows containing missing values (geom_point).
```

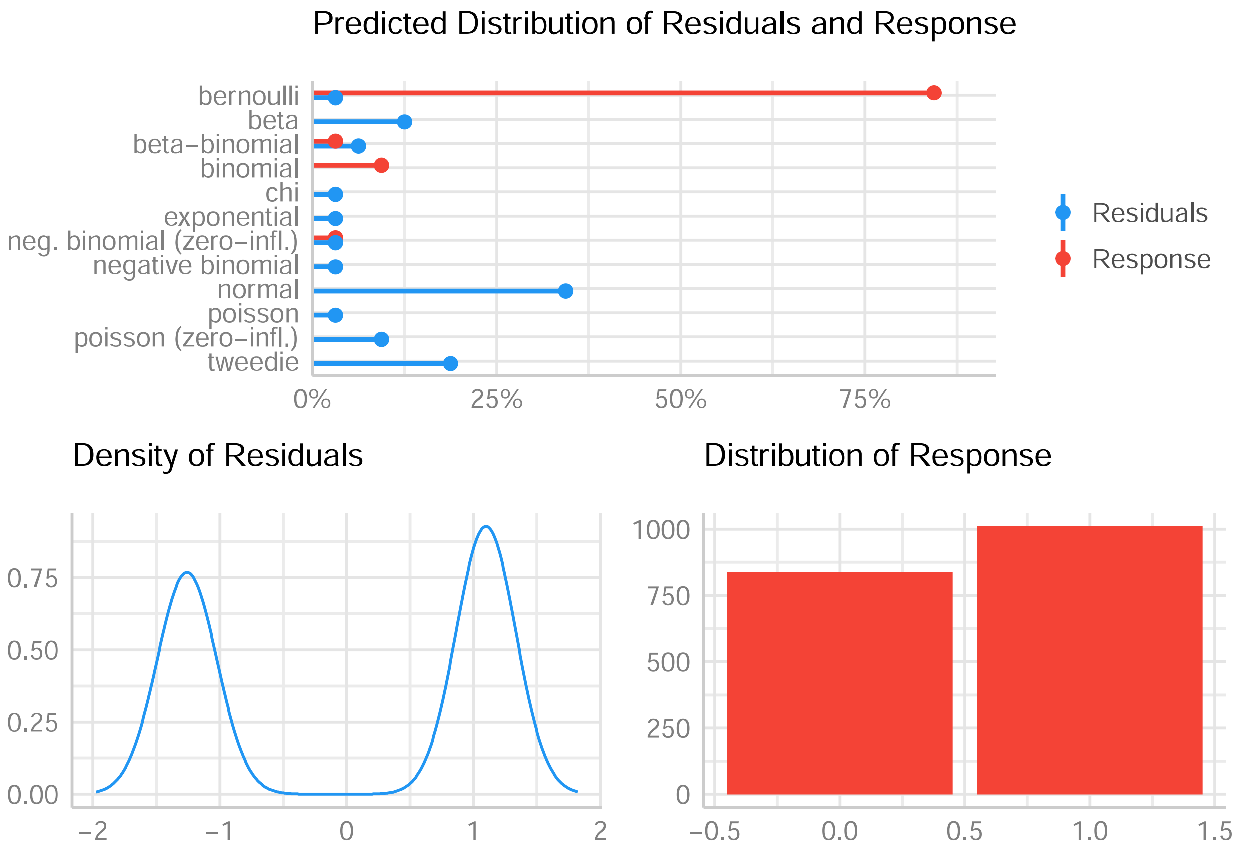

Predicted Distribution of Residuals and Response
bernoulli
beta
beta−binomial
binomial
chi
exponential
neg. binomial (zero−infl.)
negative binomial
normal
poisson
poisson (zero−infl.)
tweedie
0%
25%
50%
75%
Residuals
Response
Density of Residuals
0.75
0.50
0.25
0.00
−2
−1
0
1
2
Distribution of Response
1000
750
500
250
0
−0.5
0.0
0.5
1.0
1.5

### Fertility

#### Goal

We compared the fertility (total number of progeny obtained from a given number of females) of the SM9 line and the wild-type line.

#### Method

The effect of lines on fertility was tested using linear model and normal distribution assumptions.

#### Model output

```
##
## Call:
## lm(formula = "fertility ~ Line", data = fertil)
##
## Residuals:
##    1    2    3    4    5    6
##  489 -335 -154  563   29 -592
##
## Coefficients:
##             Estimate Std. Error t value Pr(>|t|)
## (Intercept)   1450.0      294.9   4.918  0.00794 **
## LineWT         285.0      417.0   0.683  0.53186
## ---
## Signif. codes:  0 '***' 0.001 '**' 0.01 '*' 0.05 '.' 0.1 ' ' 1
##
## Residual standard error: 510.7 on 4 degrees of freedom
## Multiple R-squared:  0.1046, Adjusted R-squared:  -0.1193
## F-statistic: 0.4671 on 1 and 4 DF,  p-value: 0.5319
```

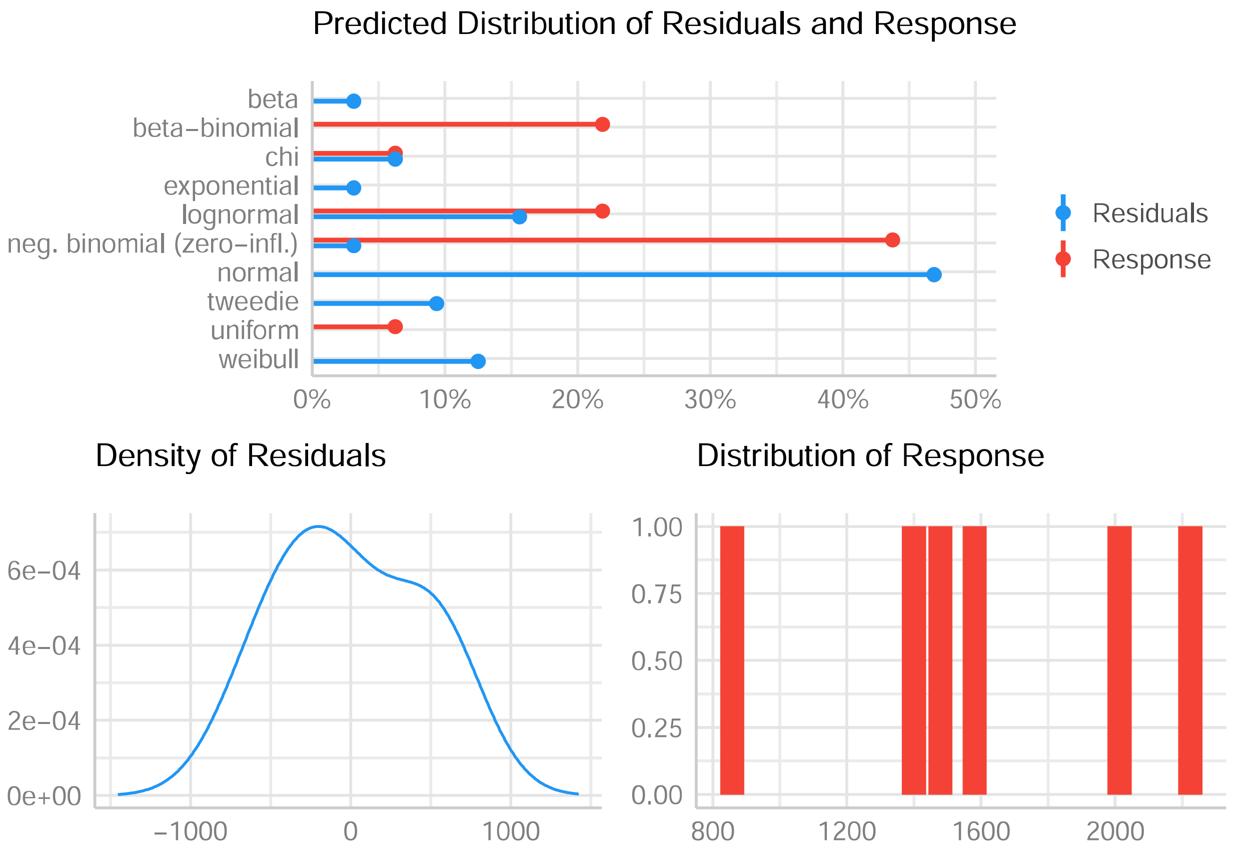

Predicted Distribution of Residuals and Response
beta
beta−binomial
chi
exponential
lognormal
neg. binomial (zero−infl.)
normal
tweedie
uniform
weibull
0%
10%
20%
30%
40%
50%
Residuals
Response
Density of Residuals
6e−04
4e−04
2e−04
0e+00
−1000
0
1000
Distribution of Response
1.00
0.75
0.50
0.25
0.00
800
1200
1600
2000

### Relative competitiveness of SM9 compared to WT

#### Goal

We measured relative competitiveness between SM9 males and wild-type males by mixing equal numbers of transgenic and wild-type males with wild-type females. The percentage of transgenic progeny was used as an indicator for relative competitiveness.

#### Method

The effect of the lines on competitiveness was tested using linear generalised model and binomial distribution assumptions. For this test, we compared the number of SM9 male progeny measured in the offspring of each competitiveness replicate, to the expected number of progeny that would have been obtained if both lines were as competitive.

#### Model output

```
##
## Call:
## glm(formula = "result ~ line", family = binomial(link = logit),
##     data = compet.Bernou)
##
## Deviance Residuals:
##     Min       1Q   Median       3Q      Max
## -0.8086  -0.8086  -0.4644  -0.4644   2.1357
##
## Coefficients:
##             Estimate Std. Error z value Pr(>|z|)
## (Intercept) -0.95011    0.02972  -31.97   <2e-16 ***
## lineobs     -1.22271    0.05307  -23.04   <2e-16 ***
## ---
## Signif. codes:  0 '***' 0.001 '**' 0.01 '*' 0.05 '.' 0.1 ' ' 1
##
## (Dispersion parameter for binomial family taken to be 1)
##
##     Null deviance: 10971  on 11264  degrees of freedom
## Residual deviance: 10383  on 11263  degrees of freedom
## AIC: 10387
##
## Number of Fisher Scoring iterations: 4
```

```
## # A tibble: 2 x 3
##   line  estimate      SE
##   <fct>    <dbl>   <dbl>
## 1 th       0.279 0.00598
## 2 obs      0.102 0.00404
```

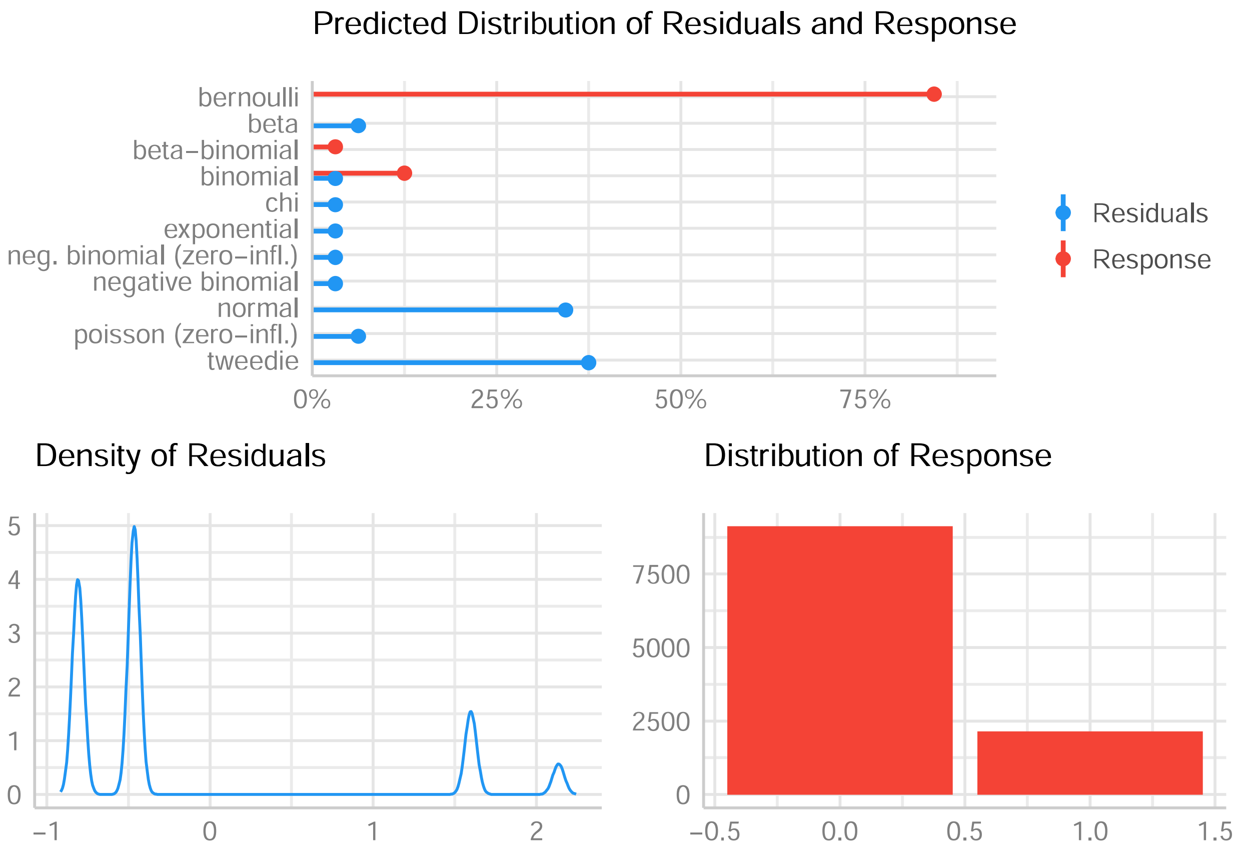
Predicted Distribution of Residuals and Response
bernoulli
beta
beta−binomial
binomial
chi
exponential
neg. binomial (zero−infl.)
negative binomial
normal
poisson (zero−infl.)
tweedie
0%
25%
50%
75%
Residuals
Response
Density of Residuals
5
4
3
2
1
0
−1
0
1
2
Distribution of Response
7500
5000
2500
0
−0.5
0.0
0.5
1.0
1.5

### Relative competitiveness of 1.2G compared to WT

#### Goal

We measured relative competitiveness between 1.2G males and wild-type males by mixing equal numbers of transgenic and wild-type males with wild-type females. The percentage of transgenic progeny was used as an indicator for relative competitiveness.

#### Method

The effect of the lines on competitiveness was tested using linear generalised model and binomial distribution assumptions. For this test, we compared the number of SM9 male progeny measured in the offspring of each competitiveness replicate, to the expected number of progeny that would have been obtained if both lines were as competitive.

#### Model output

```
##
## Call:
## glm(formula = "result ~ line", family = binomial(link = logit),
##     data = compet1.2g.Bernou)
##
## Deviance Residuals:
##     Min       1Q   Median       3Q      Max
## -0.7497  -0.7497  -0.5171  -0.5171   2.0388
##
## Coefficients:
##             Estimate Std. Error z value Pr(>|z|)
## (Intercept) -1.12547    0.01951  -57.67   <2e-16 ***
## lineobs     -0.81916    0.03200  -25.60   <2e-16 ***
## ---
## Signif. codes:  0 '***' 0.001 '**' 0.01 '*' 0.05 '.' 0.1 ' ' 1
##
## (Dispersion parameter for binomial family taken to be 1)
##
##     Null deviance: 27202  on 28395  degrees of freedom
## Residual deviance: 26516  on 28394  degrees of freedom
## AIC: 26520
##
## Number of Fisher Scoring iterations: 4
```

```
## # A tibble: 2 x 3
##   line  estimate      SE
##   <fct>    <dbl>   <dbl>
## 1 th       0.245 0.00361
## 2 obs      0.125 0.00278
```

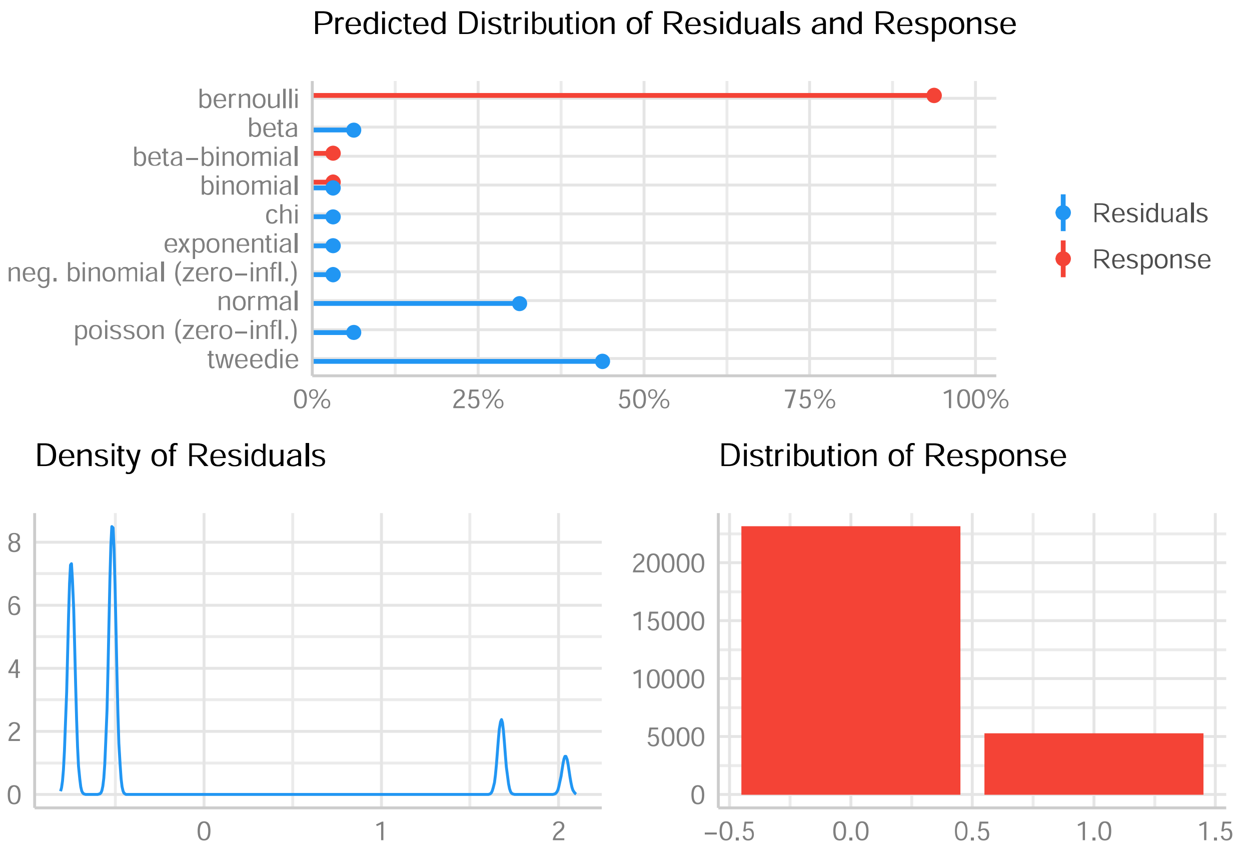
Predicted Distribution of Residuals and Response
bernoulli
beta
beta−binomial
binomial
chi
exponential
neg. binomial (zero−infl.)
normal
poisson (zero−infl.)
tweedie
0%
25%
50%
75%
100%
Residuals
Response
Density of Residuals
8
6
4
2
0
0
1
2
Distribution of Response
20000
15000
10000
5000
0
−0.5
0.0
0.5
1.0
1.5

#### Supplementary Notes

##### Supplementary Note 1: Obtaining *Nix*-expressing transgenic lines

Because additional exons were unknown when this project began, we first built pB-E1 I1 E2-OpIE2-eGFP, a transgenesis plasmid comprising only *Nix* exons 1, intron 1 and exon 2 under the control of the endogenous Nix promoter (2kb preceding the ATG start codon, **Figure S1**). We included an OpIE2-eGFP fluorescence marker for identification of transgenic individuals (the plasmid and its full sequence are available as Addgene #173505). About 300 *Ae. albopictus* embryos were injected. Among the surviving G0 adults, we obtained 24 males showing transient expression (TE) of the fluorescent marker and few females, some of which showing deformed genitalia at the pupal stage. After crossing *en masse* TE males with wild-types females, we recovered 29 eGFP positive G1 pupae, including 22 males and 7 females. We further outcrossed the 22 positive males *en masse* with wild-type females. In the G2 generation, we obtained 395 males expressing GFP while 53 were negative, and 185 females expressing GFP while 151 were negative. Of these, 12 randomly selected eGFP males were outcrossed individually with WT females, creating the lines named herein SM1 to SM12. Two crosses, (SM8 and SM10), did not produce eggs and were not further investigated. In all the other progenies but SM4, 100% of the G3 males expressed GFP. GFP positive females were also observed, although these were rare or absent in most lines. Interestingly, in SM9 57% of eGFP positive females (larger body size and female antennae) showed an intersex phenotype with deformed genitalia **(Figure S2)**. Transgenic lines were amplified over several generations and in five of them (SM1, SM3, SM7, SM9, and SM12), 100% of the males were GFP positive, suggesting either tight genetic linkage of one transgene copy to the endogenous M-locus, or transgene-mediated masculinization in the absence of endogenous M -locus.

The second injection mix comprised three different transgenesis plasmids to express the newly reported *Nix* isoforms [2], labelled with different fluorescence markers (eGFP, YFP, and DsRed, see Methods) under the control of the *Ae. aegypti* poly-ubiquitin promoter (PUb). We injected about 700 embryos with this mix, from which approximately 100 TE individuals were obtained. Similar to our previous observations, we obtained mostly male G0 TE individuals, that were split in 5 pools of males, and outcrossed *en masse* with wild-type females. Strikingly, in the G1 generation, 70-100% of the transgenic individuals were males. Some of the transgenic females in at least three pools were deformed or partially masculinised. Most transgenic G1 individuals expressed more than one fluorescent marker, indicating that they carried multiple piggyBac insertions with different transgenes. In each of 4 G1 pools, 8 transgenic males were randomly selected and outcrossed individually with WT females in order to generate lines carrying a single fluorescent marker. From these 32 individual funder males, we retained 7 independent lines in which 100% of the males were fluorescent (namely 1.2R, 1.2G, 2.2G, 3.1YR, 3.1G, 4.4Y, 5.1GR) for further analysis.

**Supplementary Note 2: Identification of *myo-sex* and *myo-fem* orthologues**

We wondered whether an essential orthologue of *myo-sex* was present in the *Ae. albopictus*'s M-locus. Using the current version of the *Ae. albopictus* genome (Vectorbase release 52, 20 May 2021), we found two homologous copies of *Ae. aegypti myo-sex* located on two distinct unplaced scaffolds (SWKY01000423 and SWKY01000369), annotated under the identifier references LOC109430926 and LOC109412105, respectively [4,5]. We designed primer pairs specific of each of these two genes, exploiting differences in their non-coding regions. Interestingly, one of these PCR markers suggested the presence of a male-specific copy of *myo-sex*, characterized by a 664 bp deletion in its non-coding sequence (**Figure S7**). These results do not align with the currently available genomic data, and suggest that a third, M-linked copy of the *myo-sex* gene exists or that the genome assembly concerning one of the above-mentioned copies is erroneous possibly due to high similarity between them.

Another sex-specifically expressed gene is *myo-fem*, described in *Ae. aegypti* as essential for female flight [6]. In *Ae. albopictus*, this gene has several potential orthologues. We compared their expression by RT-PCR between WT males and females and identified a putative *myo-fem* orthologue (LOC109402113) based on its strong female-specific expression pattern. Notably, another potential orthologue located on the same genomic scaffold as LOC109402113 and annotated as a pseudo-gene with detectable RNA expression, LOC115254984, displayed a very similar expression profile (**Supplementary Note 2, Figure 1**).

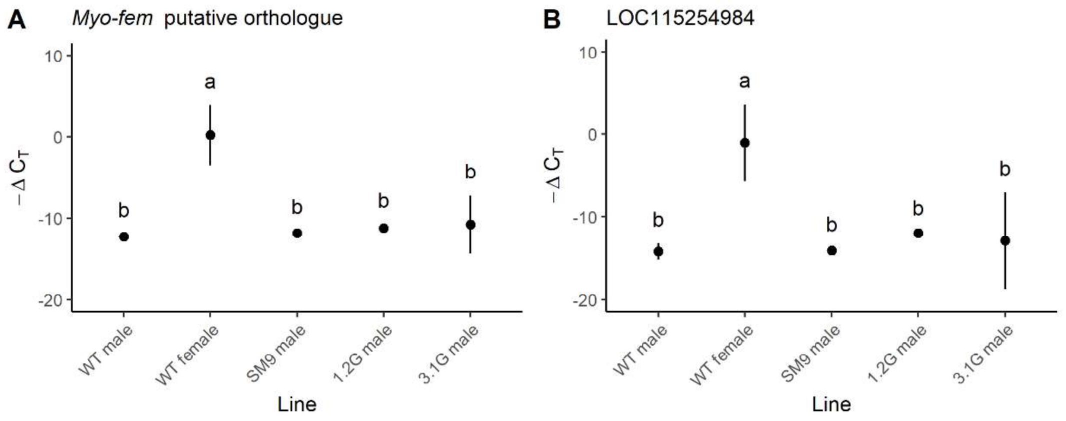


**Supplementary Note 2, Figure 1: Comparison of the expression profiles of LOC109402113 and LOC115254984, candidate *myo-fem* orthologues.**

RT-qPCR results are represented by $-\Delta C_T$, which reflects the relative expression level of each gene in a given treatment, $C_T$ values being inversely proportional to the expression levels. *AalRps7* was used as endogenous reference gene. On each panel, distinct letters represent significant difference in pairwise Tukey test ($p < 0.05$). Dots represent the mean value of the 3 biological replicates, vertical lines represent 95% confidence interval (CI). **A)** Relative expression of the candidate orthologue of the *Ae. aegypti myo-fem* gene, LOC109402113. **B)** Relative expression of the candidate orthologue LOC115254984, which is annotated as a putative pseudo-gene. This primer pair could also amplify LOC115254986 due to high sequence similarity but this other pseudo-gene seems not to be expressed.
